# INS-fMRI reveals a mesoscale limbic organization associated with medial pulvinar in primates

**DOI:** 10.64898/2026.01.05.697825

**Authors:** Yuqi Feng, An Ping, Songping Yao, Sunhang Shi, Meilan Liu, Henry C. Evrard, Jianbao Wang, Anna Wang Roe

## Abstract

The experience of emotion comprises not only the feelings (happiness, sadness), but also behavioral expression, internal awareness, and the body’s physiological response. However, it remains unclear how the brain integrates these disparate aspects of emotion. Here, to examine functional connectivity in the limbic system of macaque monkeys, we combined focal Infrared Neural Stimulation of a sensory gateway (medial pulvinar PM), with ultra-high-field 7T functional magnetic resonance imaging (INS-fMRI). We find connected sites are mesoscale (millimeter-scale) in size and arranged in patchy patterns across cingulate, insula, and amygdala. Non-overlapping connections evoked from three sequential stimulation sites in PM form clusters of multi-site integration, and appear related to known functional organization within these limbic regions. We suggest these mesoscale functional connections link the limbic axes of motor expression (cingulate), interoception (insula), and emotion-related processing (amygdala), and that, much like visual system, the limbic system is fundamentally quite orderly at mesoscale. Our results underscore the importance of millimeter-scale precision and organization in diagnosis and treatment of affective disorders.

## Introduction

Emotions are very central to human behavior; however, how emotions are represented in the brain is not well understood. Understanding the neural basis of emotion will help provide insight into human psychology and neuropsychiatric disease. Thus far, evidence from studies in both humans and animals suggest that distinct components of emotion (e.g., feelings, sensations, interoception, motor expression, and the body’s physiological response) are associated with different brain areas^1,2^. However, there is little understanding of the organization of representation within each area and how these organizations are linked between areas to generate specific emotion-related behaviors. Here, we approach this question by studying the organization of connectivity between the pulvinar (a gateway for processing sensory inputs) and major emotion-related centers in the brain. A novel method (INS-fMRI) is used to study functional connectivity *in vivo* at mesoscale.

### The medial pulvinar is a major thalamic gateway to limbic system

The pulvinar is a major thalamic nucleus which integrates multiple sensory inputs (primarily visual but also auditory and somatosensory) from the environment and disseminates this broadly to provide context for goal-directed behaviors^3,4^. One subnucleus of the pulvinar, the medial pulvinar (PM), contributes heavily to emotional processes^5,6^. In humans, (1) damage to pulvinar that includes PM is known to lead to impairments in emotional stimulus recognition^7,8^, (2) aberrant functional connectivity of pulvinar is linked to mental disorders such as major depressive disorder^9,10^, schizophrenia^11^, and generalized social anxiety disorder^12^, and (3) studies using emotional face stimuli have demonstrated PM’s modulatory effects on the amygdala^6,13^, face responses in the temporal lobe^14,15^, as well as effects in insular cortex^15^. In addition, distinct from lateral pulvinar PL and inferior pulvinar PI, PM exhibit robust anatomical connections with multiple axes of the limbic system in nonhuman primates. These include parameters of emotion-related motor expression^16–18^ (cingulate cortex^19–22)^, interoceptive self-awareness (insula cortex^21,23,24^), and sensory, cognitive, and autonomic processes (amygdala^21,25^). Functional evidence (from human and monkey studies using electrical stimulation^26,27^, resting-state connectivity^28^, task-related neuroimaging^29–31^, as well as meta-analysis of PM connectivity^32^) all demonstrate that there are prominent functional connections between PM and these three limbic areas. Together, these findings suggest that PM is a major gateway for sensory-limbic interactions^3^.

### Organizational principles of connectivity in the brain

In the primate visual system, multiple lines of evidence suggest that visual parameters in the cortex are organized at mesoscale (columnar scale)^33–37^ and are connected in a functionally specific fashion^38–42^. A first study (using focal electrical microstimulation) examining the connectivity of single columns in V2 to other columns in V1 and V2 revealed (1) column-to-column connectivity is highly matched for both functional preference (e.g., color-color, luminance-luminance between thin stripes, orientation-to-like orientation between thick stripes in V2), (2) the presence of repeating, canonical microcircuits across different stripe types, and (3) that these local micro-networks shift topographically as a whole (with shifting stimulation site)^66^. These data provide a multi-tiered organization in which visual inputs connect to: (a) specific cortical areas (e.g., V1 to V2, V3, or V4), (2) functionally specific zones within each area (e.g., thin, pale, and thick stripes in V2), and (3) within each zone, organized arrangements of functionally specific patches (e.g., hue domains within thin stripes in V2). Here, we hypothesize that functional connections between PM and limbic regions of the brain share similar organizational principles. In cingulate cortex, there are subdivisions relating to body topography (head, body and leg)^43–46^. In insular cortex, functional subdivisions relate to ‘primary-interoceptive’, ‘self-agency’, or ‘social-agency’^47,48^. In amygdala, there are subnuclei related to sensory^49,50^, cognitive^51,52^, and physiological responses^49,53^. We investigate whether, similar to the visual system, these functional subdivisions might exhibit a patterned and patchy connectivity with PM.

### A novel method for studying brain-wide mesoscale connectivity

We recently showed that mesoscale organization in the visual system extends to connections of the pulvinar, examined by a novel *in vivo* functional connectivity mapping method called INS-fMRI (Infrared Neural Stimulation with functional Magnetic Resonance Imaging)^54–58^. This method induces very brief confined thermal transients which lead to changes in membrane capacitance, resulting in neuronal action potentials^59,60^. The evoked neuronal responses lead to activations at connected sites in the brain, revealing the presence of highly specific brain-wide mesoscale networks, as observed by ultra-high field 7T fMRI. The connections activated by INS are consistent with those shown by anatomical tracing studies^57^. However, distinct from anatomy, INS induces robust activations of both monosynaptic and disynaptic connections, revealing larger brain-wide networks^57^. In a previous study, we used INS-fMRI at mesoscale sites in the pulvinar (PL and PI) and revealed that functional connections with visual cortical areas (in both ventral and dorsal pathways) are mesoscale and topographic^56^. We examine whether such principles also apply to the pulvinar-limbic system.

Here, using INS-fMRI, we find evidence suggesting an organizational architecture of connectivity between PM with cingulate, insula and amygdala. Identifying organized axes of limbic space in the brain would impact our understanding of behavior and disease^61,62^.

## Results

To investigate the mesoscale functional connectivity between medial pulvinar (PM) and emotion-related brain areas (cingulate, insula, amygdala), we used INS-fMRI to stimulate focal sites from superior to inferior in the central part of PM. Based on previous anatomical evidence^20,21^, we hypothesized that the evoked functional connections would be patchy and exhibit mesoscale topographic organization.

### INS in PM reveals functional connections

In two anesthetized macaque monkeys, an optical fiber (200 μm in diameter) was inserted into the PM, and the location of the stimulation site was confirmed using anatomical MRI image (**Fig. 1B**, white arrowhead) and functional Blood-Oxygen-Level-Dependent (BOLD) response (**Fig. 1C**, white arrowhead). The evoked responses in the cingulate, insula, and amygdala (segmented as in **Fig. 1D**; see Methods for details) revealed patchy, functionally connected sites (**Fig. 1E**, evaluated by GLM correlation with the stimulation sites, FDR-corrected P<0.005). As in our previous studies^54–58^, trains of pulsed infrared neural stimulation (**Fig. 1F**, red bars) at the stimulation site evoked BOLD signal response (**Fig. 1F**, 15 cycles shown). The BOLD signal followed a typical timecourse, with an amplitude of roughly 1-2%, peaking around 5-15 s following stimulation onset. This led to a similar BOLD response at connected sites (**Fig. 1G**, taken from yellow circled activation in **Fig. 1E**). At connected sites, the BOLD amplitude was slightly smaller (around 1%), with greater variance. Increasing the intensity of the stimulation produced a corresponding increase in response amplitude, demonstrating the intensity dependence of INS stimulation (**Fig. 1H**).

**Fig. 1.**
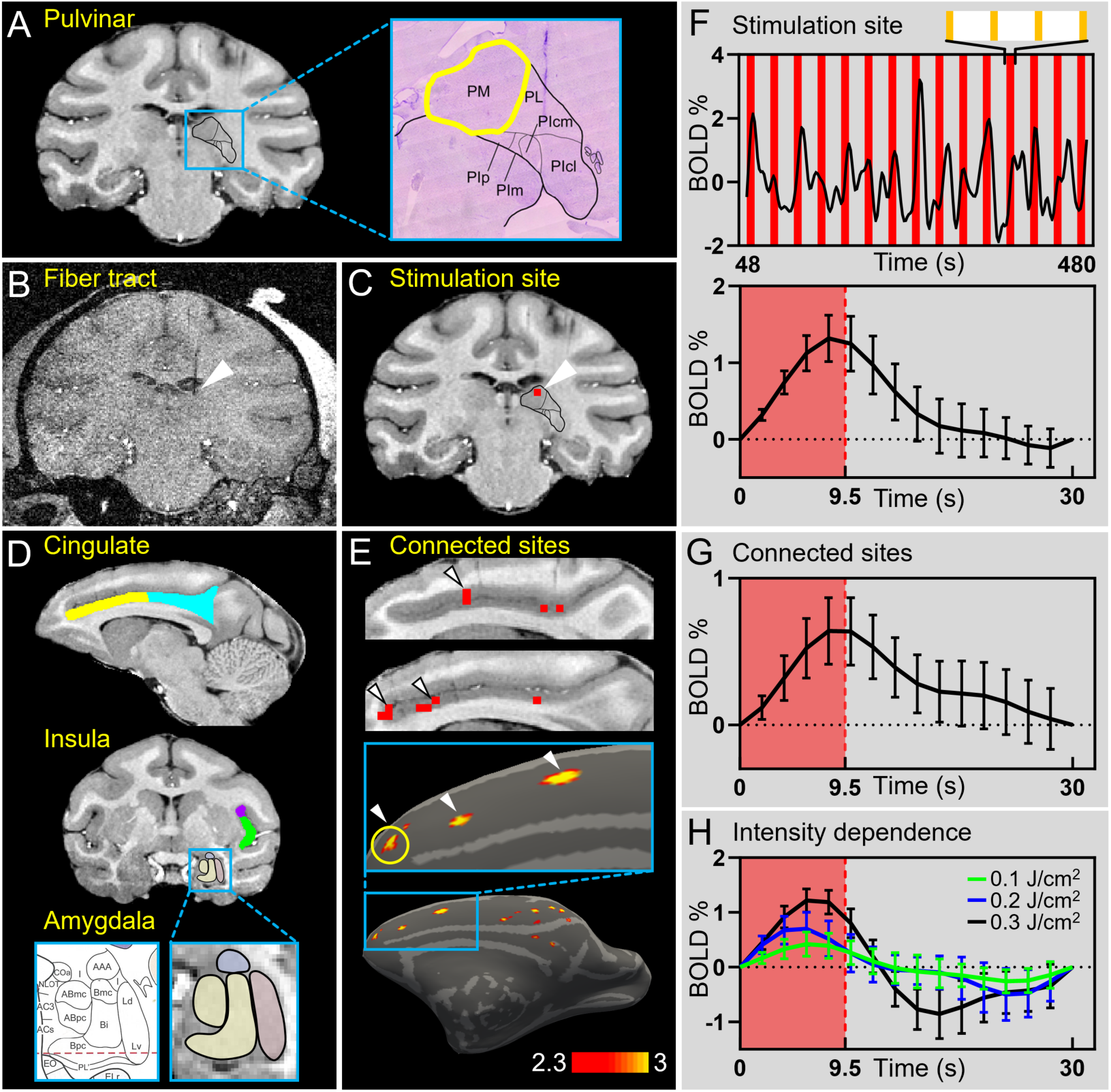
INS in PM reveals remote connections. (**A**) Parcellation of the pulvinar, based on both MRI and histological images. PM highlighted by yellow outline. (**B**) Raw anatomical image indicating the INS fiber targeting PM. White arrow: fiber tip. (**C**) White arrow: Activation at the INS fiber tip in PM (red voxel, p<0.0001). (**D**) Sample slices showing the parcellation of cingulate cortex (yellow and cyan), insula cortex (purple and green) and amygdala (zoomed-in view) in the right hemisphere, according to D99 monkey brain atlas (see Methods for details). (**E**) Above: Two adjacent slices of cingulate activations in response to INS in PM (stimulation site shown in **C,** all 3 runs combined, FDR-corrected P<0.005). Below: Surface view of the same activation. Inset: Zoomed-in view of the anterior cingulate. White arrows: corresponding positions of activated voxels seen in slice view and surface view. Yellow circle: site with timecourse shown in **H**. Color bar threshold: -log(p). (**F**) Inset above: INS paradigm consisting of four 0.5 s pulse trains (orange lines) once every 3 s. Upper graph: BOLD timecourse from the fiber tip voxel in **C**. Lower graph: Averaged timecourse from the upper graph (15 trials). (**G**) Averaged timecourse of the connected site in cingulate (yellow circle in **E,** 3 runs, 45 trials). (**H**) Example of intensity dependence at the stimulation site. The green, blue, and black BOLD timecourses show the responses at the fiber tip for INS intensities at 0.1, 0.2, and 0.3 J/cm^2^, respectively. Data are presented as mean ± SEM. Red line in graphs: 1 trial of four 0.5 s INS pulse trains. All panels are from Monkey P stimulation site P1.

### Reliability of activations revealed by INS

#### Similarity with anatomy

We compared the locations of INS-evoked patches (FDR-corrected P<0.005) with those identified in anatomical studies^21^. In Romanski’s study, WGA-HRP injections in PM revealed anterogradely labeled terminals as grey patches in cingulate, insula and amygdala. Following the tracer injection in the dorsal part of PM (**Fig. 2A**, left panel, bright label outlined by black dashed line), labeled patches were observed in cingulate area 24 and orbitofrontal areas 13 and 14 (**Fig. 2B**, left panel, red arrowheads). Following focal INS stimulation site in dorsal PM (**Fig. 2A**, right panel), functional activations were observed in similar locations in cingulate area 24, as well as areas 13l and 14c (**Fig. 2B**, right panel). The observed activation in area 45a was also observed in anatomical studies^63^. Anatomical patches were also observed in amygdala LA, BA/AB (**Fig. 2C**, left panel, red arrowheads), as well as in areas along the STS. INS activations revealed similar areas, including amygdala LA and AB, as well as STS areas TPO, PGa, and TAa (**Fig. 2C**, right panel).

**Fig. 2.**
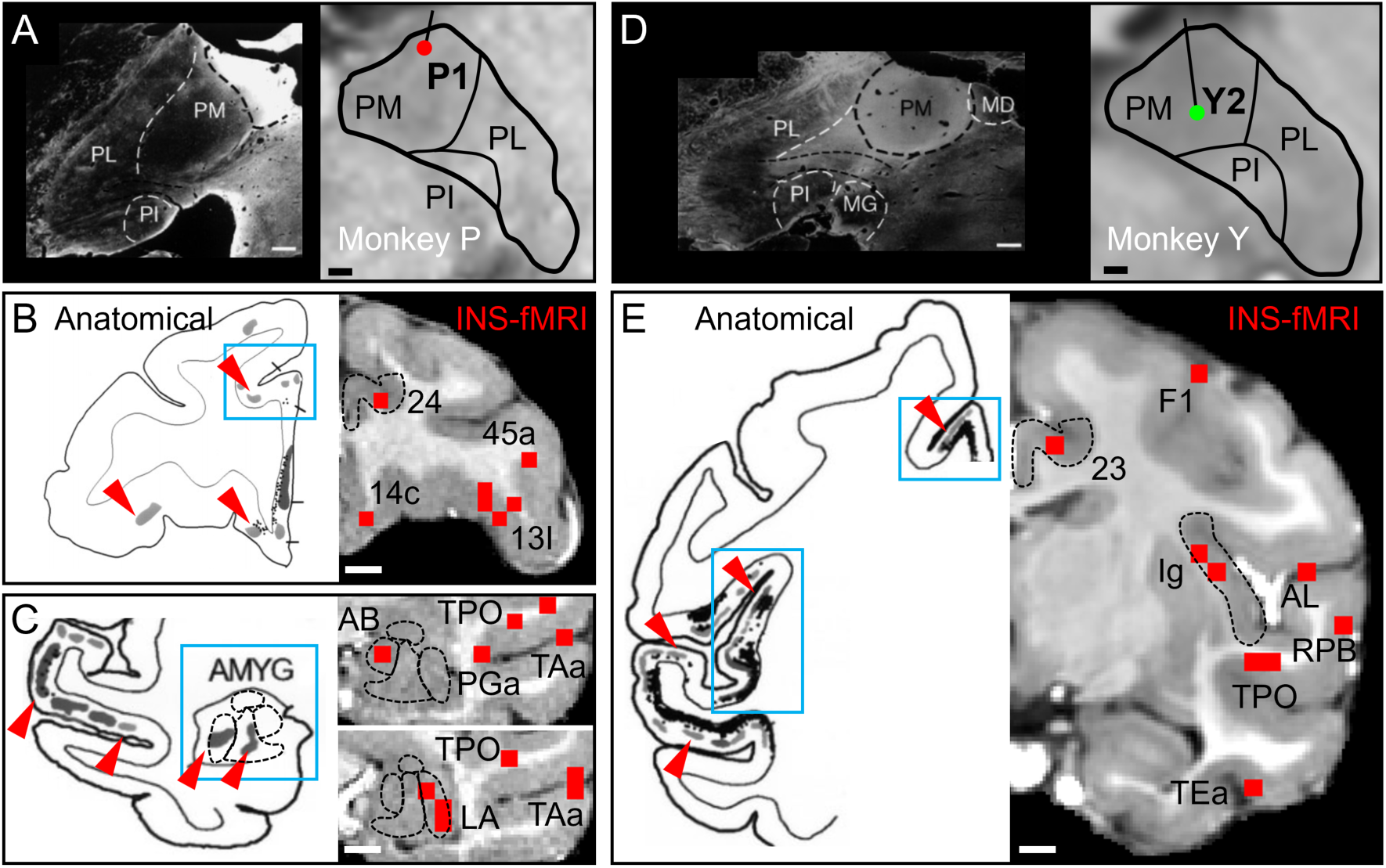
Comparison with anatomical connections. (**A-E**) Anatomical connections are derived from Romanski’s study^21^. All anatomical grey patches are from anterograde tracer WGA-HRP injection in PM, as shown in **A** and **D**. Anatomical panels are mirror symmetric to INS panels. Only ipsilateral connections are presented. (**A**) Similar location of tracer injection site in dorsal PM (left panel, from Fig. 5^21^) and INS stimulation site P1 in Monkey P (right panel). Connected sites are shown in **B** and **C**. (**B**) Left panel (from Fig. 5^21^): Anatomical grey patches in cingulate area 24, and areas 13 and 14 (red arrowheads). Right panel: Similar INS activations are observed in areas 24, 13, and 14. Activation was also observed in 45a. (**C**) Left panel (from Fig. 8^21^): Anatomical grey patches in amygdala LA, BA/AB, and STS (red arrowheads). Right panel: Similar INS activations are observed in amygdala LA and AB, as well as TPO and TAa of STS. (**D**) Similar location of tracer injection site in central PM (left panel, from Fig. 4^21^) and INS stimulation site Y2 in monkey Y (right panel). Connected sites are shown in **E**. (**E**) Left panel (from Fig. 8^21^): Anatomical grey patches in cingulate area 23, in insula Ig, as well as SII, auditory belt and STS (arrowheads). Right panel: Similar INS activations are seen in areas cingulate 23, insula Ig, auditory AL and RPB, and STS area TPO. Activations are also seen in TEa and F1. Scale bars: 800 μm in **A** and **D**, 3 mm in **B**, **C**, **E**.

We then compared a WGA-HRP injection in central PM with an INS stimulation site in ventral part of central PM (**Fig. 2D**). Anatomical patches were observed in cingulate area 23, insula, upper (SII) and lower (auditory) banks of the lateral sulcus, and STS (**Fig. 2E**, left panel, red arrowheads). INS-evoked functional activations were observed in similar locations in cingulate area 23, insula Ig, auditory belt areas AL and RPB, as well as STS area TEa and primary motor cortex F1 (**Fig. 2E**, right panel). Note that the INS-evoked connections are generally more focal, often revealing a subset of the anatomical tracer label, a result predicted by the specificity of the INS stimulation site compared with the broader anatomical injection site. There are instances of activation by INS that lacks a similar anatomical label (e.g., Tea and F1 in right panel of **Fig. 2E**), potentially reflecting a disynaptically connected activation. In addition, based on a rough estimate from the scale bar, the anatomical patches have an approximate diameter of 2 mm, which is comparable to the diameter of the INS-evoked patches (quantified in the following Results).

#### Reproducibility and stability

The reproducibility of these connections was evaluated by comparing the independent odd and even trials derived from all experimental runs (**Fig. S1**). The patterns were quite similar, as highlighted by examples of consistent activation patches across odd and even trials (white arrowheads). Additionally, we utilized another reliability test by examining activations at different thresholds. The activation patterns remained consistent across different thresholds (different p-values in **Fig. S2,** white arrowheads). Thus, INS-fMRI leads stable, reproducible, and robust BOLD responses at functionally connected locations in the brain.

### Functional connectivity in cingulate cortex

In primates, the cingulate cortex is a key brain region that processes emotions, playing a central role in integrating emotion-related information and participating in a wide range of emotion-related behaviors^17,18^, while also contributing to high-order cognitive functions such as reward processing and decision-making^64,65^. Previous anatomical studies have reported that PM connects with cingulate area 24 and area 23^19–21^. Evidence from Romanski’s study^21^ suggested the presence of patchy connections from PM to cingulate, raising questions about the mesoscale organization of functional connectivity between PM and cingulate. In addition, our previous studies in visual cortex and in amygdala showed that shifting the stimulation site leads to systematic shifting of the connected sites, revealing an organized network architecture^55,57^. To investigate these questions, we conducted sequential INS stimulation with a spacing of ∼1 mm, from superior to inferior in PM of two monkeys, with 3 sites (P1, P2, P3) in monkey P and 3 sites (Y1, Y2, Y3) in monkey Y (**Fig. 3B**, see details in **Fig. S4**). Note that the PM stimulation sites differed in two monkeys: one was primarily located in the dorsal PM (monkey Y), while the other was located more ventrally (monkey P). Our aim was not to replicate the exactly same stimulation sites across animals; rather, it was to characterize the organization of connectivity within each individual monkey. We examined the functional connections in cingulate area 24 and area 23, these two regions were defined based on the D99 atlas alignment (**Fig. 1D**, see Methods) and subsequently projected onto the reconstructed monkey brain surface (black dashed lines in **Fig. 3A**).

**Fig. 3.**
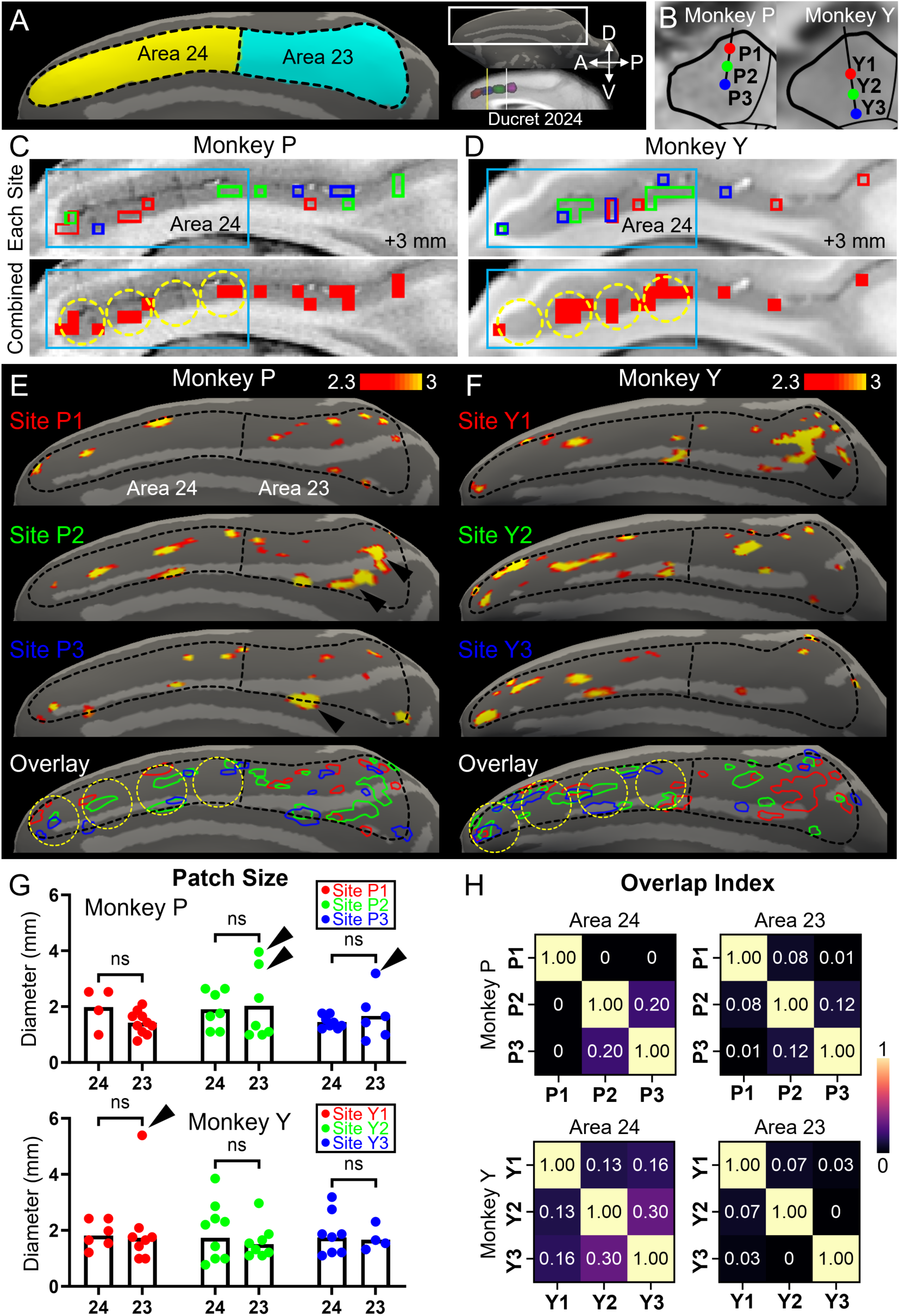
Mesoscale connected sites in cingulate. (**A**) Parcellation of cingulate cortex. Black dashed outlines indicate area 24 and area 23 of the cingulate cortex. (**B**) 3 stimulation sites in PM of monkey P and monkey Y. (**C-D**) The upper panels show the color-coded outlines of activated voxels from each PM stimulation site (FDR-corrected P< 0.005, site P1/Y1: red, site P2/Y2: green, site P3/Y3: blue) in Monkey P (**C**) and Monkey Y (**D**). The lower panels show the combined activations of the three sites in Monkey P (**C**) and Monkey Y (**D**). Four equally spaced dashed yellow circles indicate the clustering pattern in area 24 (blue box). (**E-F**) The surface view of cingulate activation maps for stimulation site P1/Y1 (row 1), site P2/Y2 (row 2), site P3/Y3 (row 3), and the overlay of these three cingulate activation maps (row 4) in PM of Monkey P (**E**, FDR-corrected P<0.005) and Monkey Y (**F**, FDR-corrected P<0.005). Four equally spaced dashed yellow circles indicate the clustering pattern in area 24, Color bar threshold: -log(p). (**G**) The diameters of activation patches in Monkey P and Monkey Y. Black arrows indicate the corresponding patches (black arrows) shown in **E** and **F**. (**H**) Dice matrices showing the patch overlap index of three stimulation in area 24 and area 23 in monkey P (P1, P2, P3) and monkey Y (Y1, Y2, Y3). Color bar: overlap index values.

We found that INS stimulation in different sites of PM led to significant activations (FDR-corrected P<0.005) in both cingulate area 23 and area 24 in both monkeys. These activations, color-coded by stimulation (sites P1 and Y1: red, sites P2 and Y2: green, sites P3 and Y3: blue), appeared as small, largely non-overlapping patches that formed semi-regular clusters along the anterior-posterior extent of the cingulate cortex (upper panels of **Figs. 3C–3D**). Following a previous study that identified three or four clusters within macaque cingulate area 23^67^, we used four equally spaced dashed circles to indicate the consistent clustering pattern of the combined activations across the three stimulation sites (lower panels of **Figs. 3C–3D**).

As shown in surface view, the distributed patterns of these activation patches, arising from each stimulation site in PM, collectively formed a functional connectivity map with the cingulate (**Figs. 3E-3F**). For example, in monkey P and stimulation site P1 (red), there were four activation patches in area 24 (mainly in dorsal part, area 24c) and 10 activation patches in area 23, with patches distributed at intervals from anterior to posterior. The reproducibility and robustness of these patches are shown in **Fig. S1A** and **Fig S2**, with largely consistent patch locations across different trials and thresholds (arrowheads).

One oft-noted commonality between cortical areas is the presence of millimeter-scale functional patches (such as ‘columns’, ‘stripes’). To investigate the size of these cingulate patches, we used a threshold which identified voxels of high statistical significance (FDR-corrected P<0.005).

Based on this threshold, we measured the diameter of each activation patch (the diameter of an equivalent circular area). As shown in **Fig. 3G** (data from different stimulation sites in two monkeys), the diameters of these patches ranged from 0.77 mm to 3.96 mm (n=42, mean ± SEM: 1.70 ± 0.11 mm) in monkey P and from 0.77 mm to 5.30 mm (n=43, mean ± SEM: 1.85 ± 0.13 mm) in monkey Y. These activation patches were focal in size and no significant difference in patch size was observed between area 24 and area 23 (compare same-colored dots in **Fig. 3G**). However, some larger patches were observed in area 23 (monkey P site P2 and monkey Y site Y3); the black arrows in **Fig. 3G** indicate the dots for specific circled patches (black arrows in **Figs. 3E-3F**). Note that these large patches may consist of a cluster of smaller patches.

As shown above, there was a distribution of patchy connections from each of the sites in PM to cingulate cortex. Then we asked: what is the relationship between the connectivity patterns evoked from these different submillimeter PM sites? To examine these questions, we color-coded the connectivity patterns from each of stimulation sites P1 and Y1 (red), P2 and Y2 (green), and P3 and Y3 (blue) in surface view, and overlaid these patterns in monkey P (**Fig. 3E**, bottom panel) and monkey Y (**Fig. 3F**, bottom panel). We predicted that if the networks from different PM sites were independent, there would be little overlap between different color-coded patches.

To quantify this, we calculated the Dice index (proportion of overlap between two color-coded patch distributions, where 0 indicates no overlap and 1 indicates 100% overlap, see Methods) between each of sites 1, 2, and 3 in two monkeys, and displayed the results as matrices in **Fig. 3H**. For example, in bottom panel of **Fig. 3E**, for stimulation site P1 and site P2 in monkey P, the red and green outlines were largely non-overlapping in both area 24 (Dice index 0) and area 23 (Dice index 0.08). For all comparisons, averaged across both area 24 and area 23, the Dice index ranged from 0 to 0.2 (n=6, mean ± SEM: 0.07 ± 0.03) in monkey P and from 0 to 0.3 (n=6, mean ± SEM: 0.12 ± 0.04) in monkey Y. Thus, the degree of overlap between these patches evoked by different stimulation sites is low, indicating these patches are largely non-overlapping and each PM stimulation site forms distinct connection patterns in cingulate cortex.

From each of the three stimulation sites in PM, we observed patches distributed across different anterior–posterior parts of both area 24 and area 23. For example, in monkey P, stimulation site P1 activated patches (red patches) in different anterior-to-posterior parts in area 24 and area 23; the same pattern was roughly observed for site P2 (green) and site P3 (blue). Additionally, we aligned the circles seen in volume view (**Figs. 3C-3D**, equally divided the area 24 into 4 circles) to the surface view; We found surprisingly that, in both monkeys, the patches from different stimulation sites were well organized and clustered into 4 groups from anterior to posterior in area 24 (yellow circles in **Figs. 3E-3F**). In comparison, when area 23 activations were displayed in surface view, such clustering was less apparent in this area.

Within each circle, the cingulate patches appeared to shift with PM stimulation sites (see red, green, and blue within each cluster). Particularly in area 24 of monkey P, the patches in in the three anterior groups shifted from superior to inferior in a red/green/blue sequence, consistent with the stimulation order. This reflects a fine-grained topographic influence of distinct PM inputs. Although in area 23 patches were not so clearly clustered, each of the anterior-to-posterior regions exhibited interleaved red/green/blue distributions. These results suggest the presence of some topographic organization in cingulate cortex. This reveals both the organization of mesoscale connectivity from PM and the presence of mesoscale functional organization within the cingulate.

In sum, we find that there are mesoscale connections between PM and cingulate. In area 24, these connections are consistent with a fine organization, forming approximately four groups arranged along the anterior-to-posterior axis. In both areas 24 and 23, the presence of distinct activation patches indicates a topographic connectivity pattern arising from sub-millimeter sized stimulation sites in PM.

### Functional connectivity and organization in insular cortex

In primates, the insular cortex plays a critical role in the emergence of interoceptive bodily, emotional feelings and cognitive functions^68^. The insular cortex is classically divided into 3 cytoarchitectonic sectors: granular insula (Ig), dysgranular insula (Id), and agranular insula (Ia) (**Fig. 4A**). Architectonic examinations reveal that each of these sectors contains several distinct, sharply delimited areas^47,69^. We further divided Ig and Id into groups englobing several areas, based on a structural and functional model^48^ (see Methods). Specifically, Ig was divided into posterior Ig (IgP) and dorsal Ig (IgD), and Id was divided into dorsal Id (IdD) and ventral Id (IdV). Together with the Ia, this yielded five insular subdivisions, which we then used to investigate the connectivity from PM (**Fig. 4A**).

**Fig. 4.**
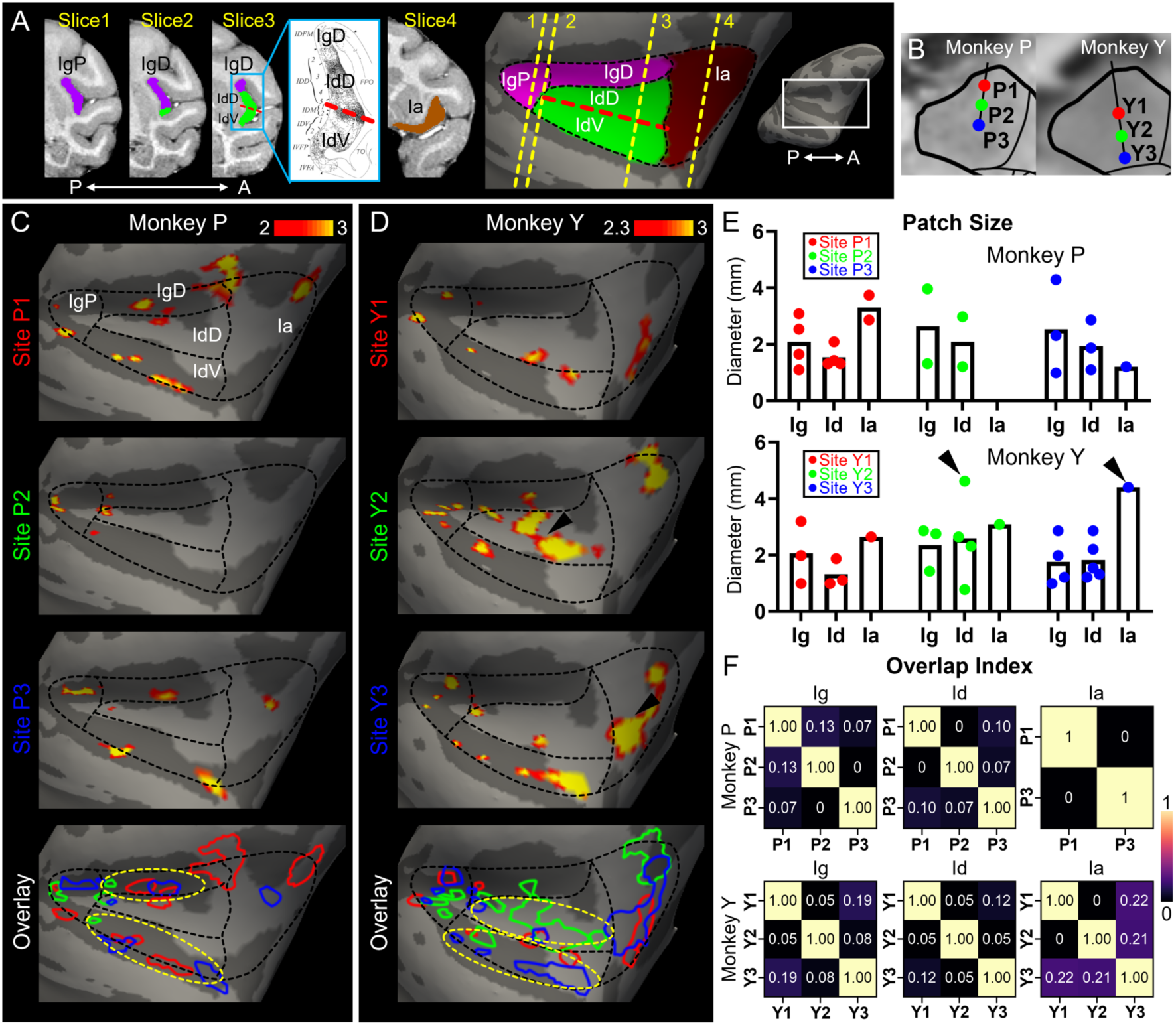
Mesoscale connected sites in insula. (**A**) Slices 1-4 show coronal MRI slices illustrating the parcellation of the insula from posterior to anterior, including granular insula (Ig, pink), dysgranular insula (Id, green) and agranular insula (Ia, brown)^48,70^. Ig was further subdivided into IgP and IgD, and Id was further subdivided into IdD and IdV (see Methods). The corresponding locations of each slice on the brain surface are shown in the right panel. (**B**) Same INS stimulation sites in PM of Monkey P and Monkey Y. (**C-D**) The surface view of insula activation maps for stimulation site P1/Y1 (row 1), site P2/Y2 (row 2), site P3/Y3 (row 3), and the overlay of these three insula activation maps (row 4) in PM of Monkey P (**C**, FDR-corrected P<0.01) and Monkey Y (**D**, FDR-corrected P<0.005). Yellow dashed ovals indicate activation patches aligned with IgD and IdV in Monkey P and with IdD and IdV in Monkey Y. Color bar threshold: -log(p). (**E**) The diameters of activation patches in Monkey P and Monkey Y. Black arrows indicate the corresponding patches (black arrows) shown in **C** and **D**. (**F**) Dice matrices showing the patch overlap index of three stimulation sites in Ig, Id and Ia in monkey P (P1, P2, P3) and monkey Y (Y1, Y2, Y3). Color bar: overlap index values.

Anatomical studies have shown that PM connects to different parts of insular cortex, including Ig, Id, and Ia^23,24^. Anterograde injections in PM revealed patchy connections in insula^21^. However, the fine-scale relationship of these connections between PM and insular had not been examined. Here, we investigated this question by examining the INS evoked activations in the subdivisions of insular cortex. We presented data from the same stimulation sites (**Fig. 4B**) and described the functional connections in five subdivisions in insula.

INS stimulation in most of the sites of PM led to significant activations in Ig, Id, and Ia (**Figs. 4C-4D**) (FDR-corrected P<0.01 in monkey P, FDR-corrected P<0.005 in monkey Y). For each stimulation site in two monkeys, these activations appeared patchy and collectively formed a functional connectivity map between PM and insular cortex. For example, in monkey P and stimulation site P1, there were 2 activation patches in the IgP, 2 patches in IgD, 4 patches in Id region (IdD and IdV), and 2 patches in Ia (**Fig. 4C)**. Based on the threshold in **Figs. 4C-4D**, we also measured the diameter of each activation patch (the diameter of an equivalent circular area). As shown in **Fig. 4E**, these activation patches were focal in size (mesoscale), with diameters ranging from 0.99 mm to 4.29 mm (n=21, mean ± SEM: 2.15 ± 0.23 mm) in Monkey P and from 0.77 mm to 4.62 mm (n=25, mean ± SEM: 2.15 ± 0.21 mm) in Monkey Y. These sizes are similar to the patch sizes observed in the cingulate (Mann–Whitney U test confirming no significant difference, **Fig. S3**). Notably, the elongated activation patches are similar in size to the ‘stripes’ labelled by tracer labelling in Id^70^. No obvious differences in patch size were observed among Ig, Id, and Ia (compare same-colored dots in **Fig. 4E**), although the small number of patches in each subregion limits a formal statistical comparison. The black arrows in **Fig. 4E** indicate the dots for specific larger circled patches (black arrows in **Figs. 4C-4D**), which may comprise a cluster of smaller patches.

To examine the relationship between these patches from different PM stimulation sites, we color-coded the connectivity patterns in insula from stimulation site P1 and Y1 (red), P2 and Y2 (green), and P3 and Y3 (blue), and overlaid these patches (monkey P (**Fig. 4C**, bottom panel) and monkey Y (**Fig. 4D**, bottom panel)). In both monkeys, we observed that the red, green, and blue outlines in Ig, Id and Ia appeared largely non-overlapping, as shown by the overlay indices in **Fig. 4F**. For example, for stimulation site P1 and site P3 in monkey P, the red and blue outlines were largely non-overlapping in both Ig (Dice index 0.07), Id (Dice index 0.10), and Ia (Dice index 0). For all comparisons, averaged across Ig, Id, and Ia, the Dice index ranged from 0 to 0.13 (n=7, mean ± SEM: 0.05 ± 0.02) in Monkey P and ranged from 0 to 0.22 (n=9, mean ± SEM: 0.11 ± 0.03) in Monkey Y. This indicates a low degree of overlap and each PM stimulation site forms distinct connection patterns in insular cortex.

We further examined the relationship between these connected patches and the dorsoventral sequence of the insula subdivisions. As shown in **Figs. 4C-4D** (bottom panel, dashed yellow ovals), the insula patches in IgD and Id (IdD and IdV) clustered into anterior-posterior elongated regions. In monkey P, patches clustered into two horizonal sulcal groups: aligning with the previously described IgD subdivision and ventral IdV subdivision. Little activation was seen in the IdD subdivision. In monkey Y, patches were observed clustered into in dorsal Id and ventral Id groups, but not in the dorsal Ig. Moreover, in two monkeys, each group contained activations (fine domains/stripes^70^) from each of the stimulation sites (red, green and blue patches), indicating a patchy influence from multiple sites in PM. Compared with this organization, the patches in posterior Ig and Ia did not show any obvious further grouping pattern. In posterior Ig and Ia, patches were also non-overlapping. However, the overlap in Ia tended to be higher, particularly in monkey Y, which may be related to the more integrative nature of Ia’s activity^71^.

In sum, we found that there are mesoscale connections between PM and insula. In dorsal Ig and Id, these fine modular connections appeared to be organized in relation to the insular subdivisions^47,48^) and to the characteristic insular stripes^70^. For all insula divisions Id, Ig, and Ia, the presence of these patches suggests that PM interfaces with all insular subdivisions.

### Functional connectivity and organization in amygdala

The primate amygdala plays a primary role in evaluating emotional salience of inputs from all sensory modalities and contributes to emotion related behaviors^50,72,73^. The amygdala consists of three primary subdivisions: CeA and AAA (central nucleus, anterior amygdala area), BA and AB (basal nucleus, accessory basal nucleus), and LA (lateral nucleus) (**Fig. 5A**, see Methods).

**Fig. 5.**
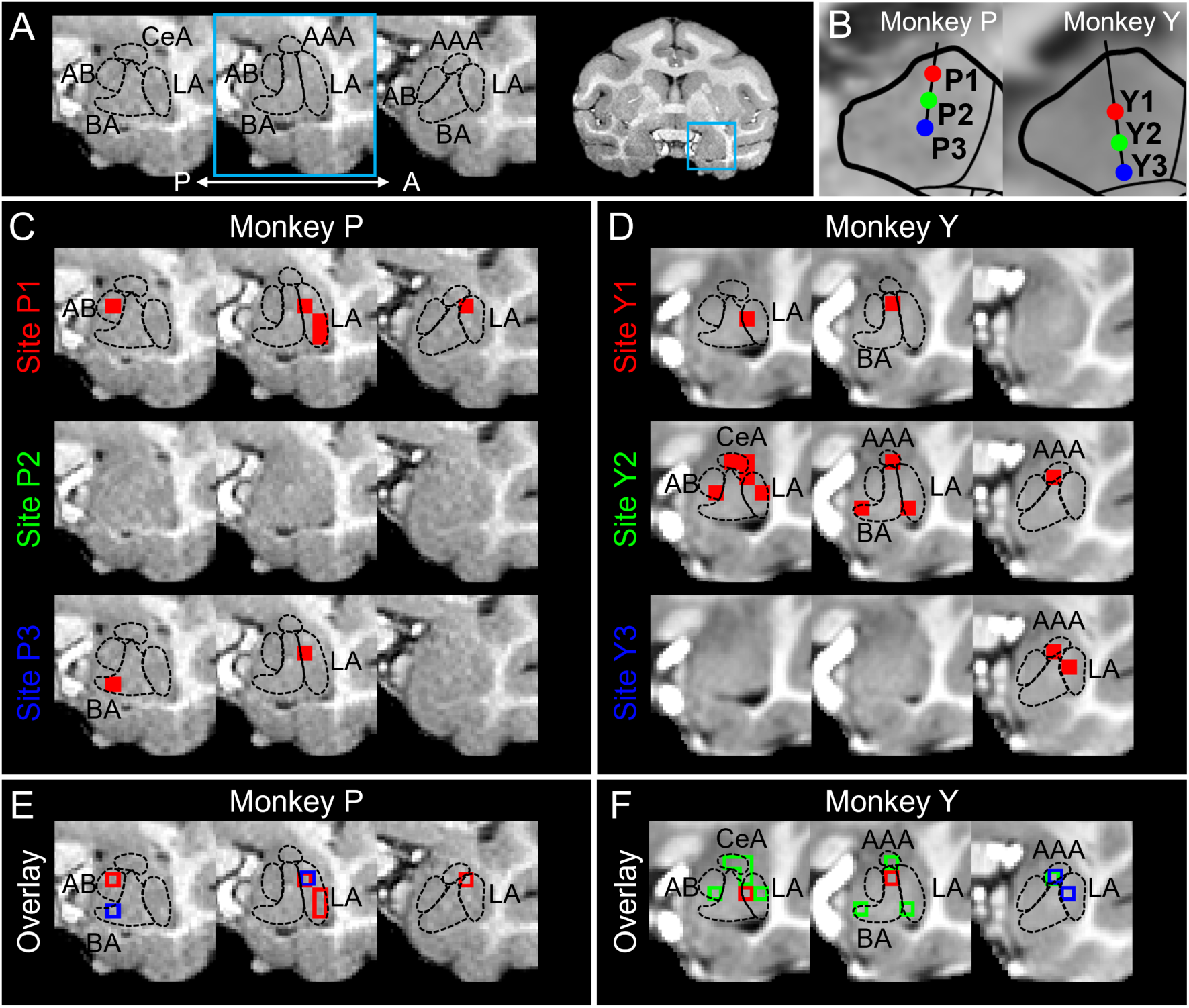
Mesoscale connected sites in amygdala. (**A**) Parcellation of amygdala. Three coronal slices are shown from posterior to anterior. Amygdala subnuclei include the accessory basal amygdala (AB), basal amygdala (BA), central amygdala (CeA), anterior amygdala area (AAA), and lateral amygdala (LA). (**B**) Same INS stimulation sites in PM of two monkeys. (**C-D**) Each row shows the amygdala activations evoked by stimulation sites P1 and Y1 (row 1), P2 and Y2 (row 2), and P3 and Y3 (row 3) in PM of Monkey P (FDR-corrected P<0.005) and Monkey Y (FDR-corrected P<0.005). (**E-F**) Overlay of amygdala activations from 3 PM stimulation sites in monkey P (**E**) and monkey Y (**F**). Color-coded outlines represent the activated voxels in **C** and **D**, evoked by PM stimulation sites P1/Y1 (red), P2/Y2 (green), and P3/Y3 (blue).

Anatomical studies have shown that PM projects primarily to the lateral of amygdala (LA)^21,25^, but the fine-scale organization of these connections remains unclear. Here, we presented data from the same stimulation sites (**Fig. 5B**) and described the functional connections between PM and the amygdala.

We identified voxels that satisfied statistical criteria (**Fig. 5C-5D**, FDR-corrected P<0.005 in two monkeys) and found that these voxels were in different subnuclei of the amygdala. For example, in monkey P and stimulation site P1 (first row of **Fig. 5C**, 3 coronal slices, from posterior to anterior, spaced 1.5 mm apart between each slice), significant voxels were seen largely in LA, with additional activation in AB. In stimulation site P2, little activation was seen (second row of **Fig. 5C**). In site P3 of monkey P and site Y1 of monkey Y, both of which were the central region of the dorsal-ventral extent of PM, voxels were seen in both BA and LA. Interestingly, voxels in CeA and AAA were only seen in the most ventral sites (sites 2 and 3 in monkey Y). Note also that site Y2 in monkey Y elicited multiple activations across all subnuclei (LA, BA/AB, CeA/AAA).

Across all stimulation sites, LA was the most commonly activated (75% in monkey P and 38% in monkey Y). We also calculated the volume of amygdala activation (see Methods). The activation volumes ranged from 3.375 mm^3^ (1 voxel) to 6.750 mm^3^ (2 voxels) (n=6, mean ± SEM: 3.938 ± 0.563 mm^3^) in monkey P and from 3.375 mm^3^ (1 voxel) to 13.500 mm^3^ (4 voxels) (n=9, mean ± SEM: 4.500 ± 1.125 mm^3^) in monkey Y. The volume of these amygdala activations was small, as 87% of activations consisted of a single voxel, consistent with the mesoscale nature of these connections (see Discussion). Overall, we revealed that PM has prominent mesoscale connections with the lateral amygdala (LA), as well as some mesoscale connections with CeA/AAA and BA/AB. Although the stimulation sites are few, this suggests that single sites in PM interface with multiple subnuclei of amygdala.

To examine whether distinct stimulation sites in PM led to distinct functional sites in amygdala, we color-coded the activations from stimulation site P1 and Y1 (red), P2 and Y2 (green), and P2 and Y2 (blue), and overlaid the outline of these voxels (monkey P (**Fig. 5E**) and monkey Y (**Fig. 5F**)). We found that these voxels were largely non-overlapping, with only one overlapped voxel in each of the two monkeys (overlapped voxels in **Figs. 5E-5F**). In sum, these distinct connections suggest the presence of functional specificity in PM-amygdala connections.

### Examination of possible topography across stimulation sites in two monkeys

In this study, we conducted 3 sequential INS stimulation sites with a spacing of ∼1 mm, from superior to inferior in PM of two monkeys. The PM stimulation sites differed between the two animals: in Monkey Y, sites were primarily located in the dorsal PM, whereas in Monkey P, sites were positioned more ventrally. Our aim was not to replicate identical stimulation sites across animals, but rather to characterize the organization of PM connectivity within each individual monkey. Given that we observed similar organizational patterns in both monkeys, we attempted to align and combine the results across the two animals, but no systematic order was found in the combined map (**Figs. S4-S6**).

This negative finding could be attributed to inter-animal variability and/or too few sites to discern topography. (1) Inter-animal variability is commonly noted in studies of electrophysiological recording and anatomical injections. Consistent with this, in visual cortical areas V1, V2, and V4, where functional maps are well established, feature maps in different animals differ in absolute stereotaxic location and even in orientation of mapping. For example, in primary visual cortex where ocular dominance (OD) columns and their eye-specific connectivity are well documented, there is much variability in the precise pattern and exact column widths across animals. (2) For the fine spacing of OD topography, due to the variability, if one were to overlay 1-mm spaced stimulation sites from OD maps in V1 in 2 animals, it would not necessarily show the expected alternating map of eye specific connectivity in V1. Likely, denser sampling of stimulation sites within each animal would be needed to examine the similarity between animals. In sum, while we found some potential distinctions between dorsal and ventral PM (Ce activations more prevalent from ventral PM sites, **Fig. S5E**), we do not have sufficient stimulation density to discern finer topography at this time.

## Discussion

Studies of both cortical^33,34^ and subcortical^74^ areas in the brain have provided evidence for submillimeter (mesoscale) functional organization. This has led to the prediction that informational networks in the brain are also composed of mesoscale nodes^54,57^. However, whether this holds true for limbic areas of the brain remains unknown. Previously, we used a novel mesoscale functional connectivity mapping method called INS-fMRI (infrared neural stimulation with functional MRI) to investigate the functional connectivity from the pulvinar (PI, PL and PM) of two macaque monkeys. The results revealed mesoscale patchy connectivity, clustered in a functional topography within the pulvinar–visual cortex networks from PL and PI; however, connectivity from PM, while similarly patchy, was largely not topographic^56^. Here, we present the functional connectivity of sequential stimulation sites in PM with limbic areas cingulate, insula, and amygdala in both monkeys. Briefly, we find evidence for patchy, topographic connectivity across these three emotion-related brain areas, suggesting potential common architecture between limbic and sensory areas.

### Methodological considerations

#### INS

INS is an optical stimulation method which is designed to produce highly focal activation around the fiber tip (∼300 μm). As a pulsed heat-based activation method, INS induces membrane capacitance change and neuronal firing^59,60^. Multiple studies have been conducted to characterize temperature rise and damage thresholds^75–77^. These studies demonstrate that this heat-mediated activation, with stimulation energies from 0.1-1.0 J/cm^2^, results in a tissue temperature rise not exceeding 2°C. For the stimulation intensities used in this study (0.1-0.3 J/cm^2^), the temperature rise does not exceed 1°C^77^ and Nissl stains of tissue in both human and nonhuman primates show these intensities are non-damaging^75,78^.

#### Mesoscale

The unique aspect of INS is its ability to achieve highly focal stimulation (200 um fiber, ∼300 um target area, submillimeter scale), which, as shown by the data, leads to focal activations in distant target regions, resulting in brain-wide columnar networks^54^. Note that, although the resolution of functional images in this study is 1.5 mm isotropic (millimeter scale), INS (of identical stimulation parameters) actually evokes a submillimeter scale response at the fiber tip and connected sites, as shown by electrophysiological^66,79,80^, optical imaging^79,81^, fMRI^54–57^, and 2-photon methods^82^. In this study, we observed a range of activation sizes (1 mm to 5 mm in diameter), with the majority of activations (75% in cingulate, 58% in insula, and single-voxel activations in amygdala) being nominally around 2 mm. We note that fMRI and optical imaging studies in other areas have shown that such 2-3 mm millimeter-scale patches contain local networks of ∼0.3-0.5 mm column-sized patches (i.e. single site stimulation evokes a local network of functionally specific patchy activation spanning 2-3mm)^83^, raising the possibility that the patch sizes measured here could be an overestimate. Conceptually, we associate ‘mesoscale functional connectivity’ with brain-wide networks of functionally connected columns evoked by submillimeter focal stimulation.

#### Advantages

INS used with these parameters is non-damaging^75,78^, and can be delivered at multiple sites of stimulation within single subjects within single sessions and in separate sessions over time. This more easily permits the study of questions that previously required injection of multiple tracers and animal sacrifice. Connectivity mapping coupled with the brain-wide field of view offered by INS-fMRI permits us to investigate the topography of connectivity, functional specificity of connections, and organization of global brain-wide mesoscale circuits^55,57^. The precision of targeting is also an advantage as desired sites can be acquired with <500um accuracy (depending on resolution of structural images). We also note that single-site stimulation leads to statistically significant activations at both monosynaptic and disynaptic connected sites, providing a broader view of networks compared to standard anatomical techniques. Furthermore, INS-fMRI is an *in vivo* technique, enabling multiple sessions of data acquisition across time (longitudinal studies). Importantly, for future clinical application, INS provides a way to study fine-scale and brain-wide functional networks without the need for prior viral transfection^62,78^. INS can also be combined with other methodologies (such as optogenetics, DREADDs, electrophysiology, behavior) for more in-depth characterization.

#### Comparison with other PM stimulation studies

Intracranial EEG studies in humans have stimulated PM and examined its connectivity with the cingulate and insular cortices^26,84^. However, because electrode implantation inherently provides limited and sparse spatial coverage, these approaches are less suited for systematically characterizing the fine-grained topographic organization of PM-related pathways. Electrical stimulation combined with fMRI (es-fMRI) has similarly revealed PM connectivity with the insula and cingulate in monkeys^27^. But, due to current spread during electrical stimulation, the resulting activations reflect the connectivity of the broader PM region rather than that of focal, submillimeter-scale sites. As a result, these stimulation methods face challenges in revealing the mesoscale topography of PM connectivity.

### Functional inferences of PM-Limbic mesoscale organization

Here, we discuss the possible relationships of our findings with known functional organizations in these areas (**Fig. 6A**). While we do not directly localize specific functions within the cingulate cortex, insula, or amygdala, we interpret the observed connectivity patterns in the context of previously published literature describing the known functional roles of these regions. Our findings show that the PM distributes to different subdivisions within each target area and that each subnucleus integrates across multiple PM sites. From this perspective, we suggest that PM participates in the integration of distinct emotion-related functions through its organized connectivity with these areas. Further studies need to be done to directly examine the functional meaning of these connections and to better understand how the PM serves as a modulator in emotion-related and limbic processing.

**Fig. 6.**
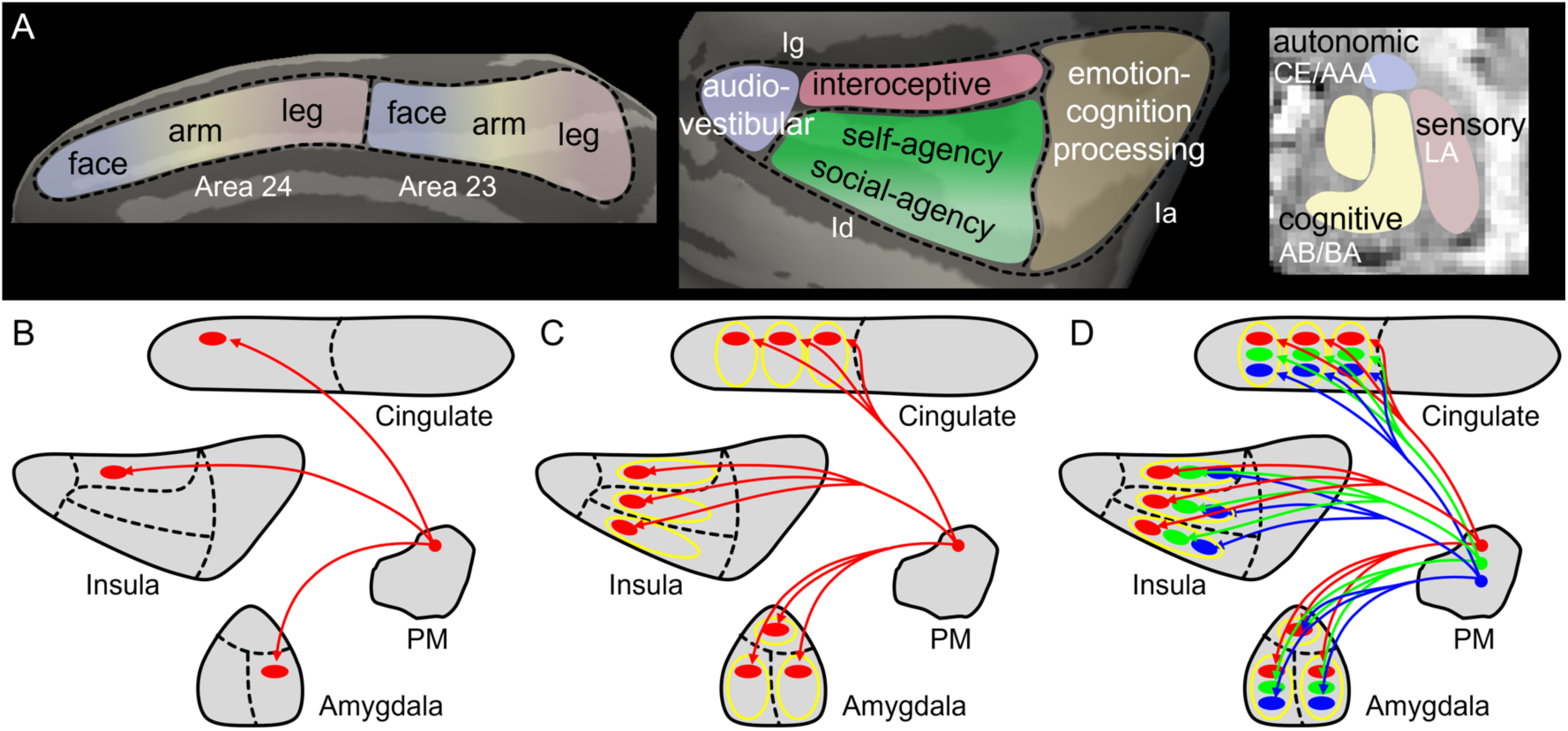
Schematic diagram of 3-tiered limbic circuit integration. (**A**) Sub-functionality interpretation of cingulate, insula and amygdala. (**B**) Single sites in PM leads to concurrent activations in cingulate, insula, and amygdala. (**C**) Single sites in PM leads to concurrent activations across multiple modalities (yellow ovals) in cingulate (face-arm-leg motor maps), insula (‘primary-interoceptive’, ‘self-agency’, and ‘social-agency’ functional subdivisions) and amygdala (autonomic, cognitive, and sensory aspects of emotion). (**D**) The sequential sites in PM connect to sequences of non-overlapping patches within each modality in cingulate (area 24), insula (Id and dorsal Ig), and amygdala (mainly LA and BA).

#### Cingulate

Here, we examined activations in areas 23 and 24 of the cingulate evoked by INS in PM. Located on the medial wall of the hemispheres, the regions falling on the gyral surface are 23a, 23b and 24a, 24b, while the portion within the cingulate sulcus is 23c and 24c. According to anatomical^44,85^ and microstimulation studies^43^, motor maps have been associated with 23c and 24c, and are reported to be organized in a face/arm/leg order from anterior-posterior in each of 23c and 24c^45^, with possible increased cortical magnification for face^67^. Human neuroimaging studies of cingulate also reported 4 clusters of somatotopic eye/tongue/hand/food movements^46^, further suggesting somatotopic similarity with monkeys, although with an increased number of clusters. Given this functional topography, we evaluated the possibility that the INS-evoked activation patches were related to the known topographic organization (**Fig. 6A**, left panel).

We found that both area 23 and 24 received extensive patchy connections, consistent with previous studies^21^, and that these connections were primarily located within the cingulate sulcus. These patches were distributed across the anterior-to-posterior extent of each area. In area 24, patches evoked from distinct PM sites (**Figs. 3E-3F**, site P1 and Y1 red, site P2 and Y2 green, site P3 and Y3 blue) exhibited a clustering pattern (yellow circles), with approximately four clusters observed along the anterior–posterior axis corresponding to the motor map. These clusters could correspond to previously described body maps (e.g., eye/face, arm, leg). In contrast, area 23 displayed less clear clustering, supporting a potentially more complex body map representation^43,44,46^ related to its role in spatial navigation^16^. Note that although somatotopy has been described in the cingulate, the essential function may be less about motor movement than about translating internal body states (related to emotional experience^86^, motivation^87^, choice^64,65^, or environmental context) into action^17^. In sum, our data suggest the presence of mesoscale, patchy connectivity between PM and cingulate, and a topographically organized clustering of multi-site PM integration. This organization may suggest that PM plays an integrative role in coordinating cingulate-related emotion functions arising from different body regions.

#### Insula

In insular cortex, multiple lines of evidence from anatomical^47,70^, functional imaging^88,89^, and behavioral studies^90,91^ supports functional specialization across insula subdivisions.

Specifically, according to the insular structural and functional model: (1) the posterior (IgP, granular) is associated with sensory (audio-vestibular), (2) the anterior (Ia, agranular) with emotion/cognition, and (3) the central (dorsal IgD and, Id, dysgranular), which contains three dorsal-to-ventral striped divisions, is associated with ‘primary-interoceptive’ (IgD, dorsal Ig), ‘self-agency’ (IdD, dorsal Id) and ‘social-agency’ (IdV, ventral Id), respectively^47,48,70^ (**Fig. 6A**, middle panel). Our data revealed mesoscale connections organized with respect to these insular functional subdivisions. In Monkey P, INS evoked patches are clustered in the ‘primary-interoceptive’ dorsal fundus of IgD and ventral ‘social-agency’ IdV^47,48,70^, while, in Monkey Y, they were clustered only in dorsal ‘self-agency’ IdD and ventral ‘social-agency’ IdV; this difference could be due either to the differences in stimulation sites or to inter-animal variability. In posterior Ig, multiple patches were observed, in each of dorsal and ventral portions of lgP, raising possible presence of distinct clusters. Some of the largest patches in the insula were observed in anterior Ia, perhaps suggesting broader integration fields. Overall, similar to the cingulate, within each functional subdivision (‘audio-vestibular’, ‘primary-interoceptive’, ‘self-agency’, ‘social-agency’, and ‘emotion-cognitive processing’), there are non-overlapping patchy connections from different PM sites, suggesting the presence of higher order interoceptive integrations from PM.

#### Amygdala

In amygdala, studies of anatomical connections suggest functional distinctions among its subnuclei: CeA/AAA, which connects to the basal forebrain, hypothalamus, and brainstem, is broadly associated with autonomic circuits^92^; the BA/AB, connected with frontal, insula, and cingulate regions, is related to cognitive circuits^93,94^; and the LA, receiving visual, auditory, and somatosensory inputs from the thalamus and cortex, is linked to sensory circuits^95^ (**Fig. 6A**, right panel). Our recent study using INS to stimulate sites in LA, BA, and Ce reveal organized patchy connectivity across sensory, motor, cognitive, associational, limbic, and autonomic areas, suggesting broad, mesoscale-based amygdala influence on integrated behavioral function^57^.

Here, we observe that, similar to the cingulate and insula, stimulation at nearly all six sites (with the exception of site P2 in Monkey P) across the two monkeys elicited activations in LA, consistent with previous anatomical studies^25^. As well, most sites exhibited additional activations in other subnuclei (such as Ce/AAA, BA/AB), indicating broad functional interactions with autonomic, cognitive, sensory subregions of the amygdala. In each subregion, particularly in LA and BA/AB (with Ce/AAA being too small for clear analysis), the lack of overlap between voxels arising from different PM inputs (see **Figs. 5E-5F**) suggests potential functional distinctions between the activated sites in amygdala. Thus, although the amygdala is a subcortical than cortical brain region like the cingulate and insula, we similarly find the presence of mesoscale organization^57,72,96,97^ in PM-amygdala connections, along with the potential coordination of different functional modalities within the amygdala by PM inputs.

### Global-to-Local scales of limbic integration by PM in circuits

We suggest that there are commonalities in the organization of mesoscale connections between PM and cingulate, insula, amygdala, reflecting multiple scales (from global to local) of emotion circuit integration, as shown in **Figs. 6B-6D**: (1) Single sites in PM led to concurrent global activations in cingulate, insula, and amygdala (**Fig. 6B**); (2) Single sites in PM led to concurrent activation in multiple modalities (functional subdivisions) in the cingulate (anterior-to-posterior body topography subdivisions, yellow circles), insula (primary-interoceptive IgD, self-agency IdD, social-agency IdV subdivisions, yellow circles), and amygdala (cognitive BA/AB, autonomic Ce/AAA and sensory LA, yellow circles) (**Fig. 6C**); (3) Sequential sites in PM (red/green/blue) connect to sequences of non-overlapping patches within each modality in cingulate (red/green/blue in each anterior-posterior region), each modality in insula (red/green/blue in IgD, IdD, IdD) and potentially different modalities in amygdala (red/green/blue mainly in LA and BA), revealing topographic connectivity relationships with each limbic area (**Fig. 6D**).

Together, this presents a concept of multi-tiered organization of functional emotion circuit integration mediated by PM. Importantly, we show that single sites in PM tie together sets of nodes across sites associated with different emotion-related modalities (**Fig. 6C**), and suggest the integration of multiple PM site inputs within local modalities/clusters (clearly seen in cingulate and insula) (**Fig. 6D**). Thus, our findings reveal the presence of organized circuitry that could underly the integration of different components of pulvinar-guided emotion-related response in the brain.

### Mesoscale architecture underlying emotion circuits

Emotion has traditionally been characterized along two dimensions: valence (ranging from positive to negative) and arousal (the intensity of emotion)^98,99^. Recent studies have suggested that these dimensional aspects of emotion are represented similarly in both behavior and the brain, with distinct topographical organizations^100^. Furthermore, other brain regions, such as the cerebellum^101^, have also been implicated in emotion processing and exhibit functional topography. Together, our findings strengthen and add a mesoscale understanding to organized representations of emotions in the brain.

In sum, the circuitry described here reveals a highly modular organization underlying how broad sensory information from the pulvinar (related to PM) orchestrates influence on distinct aspects (axes) of limbic behavior. We suggest that this is achieved via selective and integrated connections for motor behavior (different body parts) underlying emotion expression; for different aspects of internal self-awareness (including interoception, self-knowledge, and awareness of others); and for generation of sensory, cognitive, and autonomic aspects of emotion. The view that emotion-related behaviors are based in an elegant brain architecture raises the possibility that emotion-related components are fundamentally quite orderly. This perspective may provide insight into how abstract aspects of behavior may be mapped in the brain as well as impact neuropsychiatric interventions^61,62^.

## Acknowledgments

This work was supported by STI 2030 Major Projects 2021ZD0200401 (to A.W.R.); the National Natural Science Foundation of China U20A2022 and 81961128029 (to A.W.R.), 82501864 (to J.B.W), 32271049 (to H.C.E); Fundamental Research Funds for the Central Universities 2023ZFJH01-01 (to A.W.R); Shanghai Municipal Science & Technology Major Project 2019SHZDZX02 (to H.C.E); Belgian Fonds Wetenschappelijk Onderzoek-Vlaanderen G0E0520N (to H.C.E).

We thank Zhejiang University 7T Magnetic Resonance Imaging Platform and Mr. Bin Xu for assistance with MRI scanner operation and equipment coordination. We thank Lili Zhang, Shuangshuang Qian and Jun Mao from the Animal Core Facilities of Zhejiang University School of Medicine for their technical support. We thank Lixia Gao for her insightful review of the manuscript. We also thank Meizhen Qian, Feiyan Tian, Yipeng Liu, Meixuan Chen, Xiao Du, Qiuying Zhou, Lihui Li, Libo Lin, Zeng Pan, Jianfeng Fu for their assistance with experimental data collection.

## Author Contributions

Yuqi Feng: Conducted data analysis, made figures, wrote and revised paper, conducted scans, developed data acquisition and analysis methodology.

An Ping: Performed surgeries, developed data acquisition and analysis methodology. Songping Yao: Performed surgeries, developed data acquisition methodology.

Sunhang Shi: Conducted scans, developed data acquisition and analysis methodology. Meilan Liu: Performed surgeries, developed data acquisition methodology.

Henry C. Evrard: Supervised project, made figures, wrote and reviewed paper. Jianbao Wang: Supervised project, developed methodology, wrote and revised paper.

Anna Wang Roe: Supervised project, designed experiments, developed methodology, wrote and revised paper.

## Methods

### Macaque monkeys

Two hemispheres in two adult female rhesus monkeys were used in this study (Macaca mulatta; 5.5 kg for Monkey P and 5.1 kg for Monkey Y; for both two monkeys, PM in right hemisphere was stimulated (**Fig. 1A**)). We analyzed and presented 3 stimulation sites in Monkey P and 3 stimulation sites in Monkey Y (**Fig. 3B**). The data presented are part of a broader dataset acquired from PL, PI, and PM in these two monkeys^56^.

### Animal surgical procedures and anesthesia maintenance

All procedures were approved by the Institutional Animal Care and Use Committee (IACUC) of Zhejiang University, in accordance with the National Institutes of Health (NIH) guidelines.

Monkeys were placed in an MRI-compatible stereotaxic instrument after injection of ketamine (5–10 mg/kg) and atropine (0.03 mg/kg). During surgery, monkeys were artificially ventilated and anesthetized with 1-2% isoflurane. A single burr hole was drilled on the skull after confirming the coordinates of PM stimulation sites with guidance of pre-scanned high resolution MRI anatomical images and the D99 monkey brain atlas^102^. During MRI data acquisition, monkeys were artificially ventilated and anesthetized with sufentanil (induction 0.5 μg/kg, maintenance 1–2 μg/kg/h) and supplemented with 0.2-0.5% isoflurane. A light dose of vecuronium bromide (induction 0.25 mg/kg, maintenance 0.05–0.10 mg/kg/h, IV) was used to stabilize the eyes. During the entire surgery and MR scanning procedure, the animal’s body temperature was maintained at 37.5-38.5 °C with a water blanket and we kept monitoring vital signs (heart rate, SpO2, end-tidal CO2, respiration rate, temperature) of the monkeys to make sure that the anesthesia was stably maintained. Visual gratings were first presented to check activation in the visual cortex, and the monkey’s eyes were then closed during INS data acquisition. After completion of the scanning session, the animal was recovered. Antibiotics were administered daily for 3–5 days to prevent infection. Additional sites of stimulation were obtained in the same animal after at least 7-10 days of rest. Each animal underwent several anesthesia and data collection sessions.

### INS fiber preparation

We used 200 μm diameter low-OH silica core fibers (0.22 numerical aperture) with 10 μm thick cladding of fluorine-doped silica, resulting in a total diameter (outer diameter) of 220 μm. Prior to the experiment, the optical fibers were inspected, cleaned and sterilized. Using a simple inverted microscope and appropriate optical mounting hardware, both the ferrule end and the free end of the fiber probes (the latter directly contacts the brain tissue) were inspected. To remove any particles from the core and cladding at the free end, the fiber may be drawn multiple times through a folded lens tissue wetted with isopropyl alcohol. Both ends must be free of contaminants such as dust or oil to achieve optimum transmission of the laser light. The dura was punctured with a fine needle prior to fiber insertion, and the optical fiber was then inserted into through the burr hole at a predetermined angle and depth using an MR-compatible micromanipulator.

### INS paradigm

To determine the precise location of the stimulation sites within PM, we conducted a raw anatomical MRI scan for every stimulation site, which revealed a dark site of signal dropout distinct from surrounding tissues (white arrow in **Fig. 1B**, also see **Fig. S4**). Stimulation sites were also further confirmed by focal strong Blood-Oxygen-Level-Dependent (BOLD) activation around the fiber tip (white arrow in **Fig. 1C**). As in our previous studies^54–58^, a block design INS paradigm was used for neural stimulation. Each trial period of 30 s included Laser-On for 9.5 s and Laser-Off for 20.5 s; four pulse trains (1875 nm wavelength, 0.5 s per pulse train, each pulse 0.25 ms, 200 Hz) were transmitted through an optical fiber (200 μm in diameter) during the Laser-On period. These stimulation paradigms have been validated as non-damaging in our previous work^75,77,78^. Overall, in one experimental run, there were 15 trials with a total duration of 480 s (**Fig. 1F**). The intensity of laser pulses was between 0.1 to 0.3 J/cm^2^, which we has been shown to be non-damaging to brain tissue^75,77,78^. In this study, after testing intensity dependence, we report all experimental runs with intensity of 0.3 J/cm^2^.

### MRI data acquisition

The MRI data were acquired on a 7T MAGNETOM scanner (Siemens Healthineers) equipped with a SC72 body gradient (70 mT/m maximum gradient strength and 200 T/m/s maximum slew rate). A single loop RF coil with 7 cm inner and 14 cm outer diameters (RAPID MR International, Columbus, OH) was used for transmission and signal reception. Anatomical images were collected with 0.3 mm isotropic resolution using a T1-weighted MPRAGE sequence following protocol parameters: echo time (TE) = 2.82 ms, repetition time (TR) = 2590 ms, inversion time (TI) =1500 ms, flip angle = 11°, matrix size = 320 × 320, slice number = 240, bandwidth = 300 Hz/Px; BOLD functional Magnetic Resonance Imaging (fMRI) images were acquired with 1.5 mm isotropic resolution using an gradient echo EPI (Echo-Planar Imaging) sequence following protocol parameters: TE = 22 ms, TR = 2000 ms, flip angle = 90°, matrix size = 64 × 64, slice number = 35, bandwidth= 1776 Hz/Px. In monkey P, 3, 2 and 3 runs, in which 45, 30 and 45 trials for each site (P1-P3) were collected. In monkey Y, 2, 2 and 3 runs, in which 30, 30 and 45 trials for each site (Y1-Y3) were collected.

### Anatomical MRI data processing and surface reconstruction

Both MRI anatomical and functional data were analyzed using Freesurfer (version 7.1.1)^103^, AFNI (version 19.1.25)^104^, and ANTs (version 2.4.1). The anatomical images from 15-20 scans were B1 bias corrected, aligned using rigid body alignment and averaged to improve Signal-to-Noise Ratio (SNR). To reveal the topographic organization of connections in the cingulate and insular cortex, cortical surfaces were reconstructed from the high SNR anatomical image following the Freesurfer surface reconstruction workflow customized for macaque monkey at 7T MRI^56,105^: (1) Based on the high SNR anatomical image, we first adjusted the header file orientation information and voxel size information to conform to the common data processing software standards for the human brain; (2) Signal inhomogeneity was then corrected, and images were further normalized in the macaque standard space; (3) Gray matter, white matter, and cerebrospinal fluid were segmented, and the extracerebral tissue were removed; (4) Based on the segmentation results, the white-matter/gray-matter border (WM surface) and the gray-matter/CSF border (pial surface) were generated; (5) The cortical surfaces were subsequently smoothed, topology-corrected, and inflated. To better understand the morphometry of the functional organization in cortical areas spanning gyral and sulcal regions, we transformed the functional data (activations in cingulate and insula) from volume space to surface space. The statistical results in volume space were projected along the topology-corrected gray-matter/white-matter surface to avoid the pial vessel effects, using transformation estimated from BBR, with ‘cubicbspline’ interpolation. We confirmed the corresponding relationship between volume and surface views (**Fig. 1E**).

### Segmentation of cingulate, insula, and amygdala subdivisions

The D99 monkey brain atlas (version 2.0)^102^ was nonlinearly aligned to the high SNR anatomical image for brain region segmentation based on *3dQwarp* tool in AFNI. For the cingulate cortex, we segmented area 23 (cyan region, including subdivisions 23a, 23b, and 23c) and area 24 (yellow region, including 24a, 24b, and 24c). For insula, we segmented granular (Ig, purple region), dysgranular (Id, green region) and agranular (Ia, brown region), and each of these sectors can be further divided into several distinct sharply delimited areas^47^. Here, we provide further details on how the insula was subdivided into five subdivisions based on our previously defined cytoarchitectonic insular subdivisions^48,70^ (blue boxed zoom-in view in **Fig. 4A**). In insula, Ig contains two main sets of areas: the dorsal and ventral areas located in the posterior end of the insula (Igd and Igv) and the posterior, middle and anterior fundal areas (Idfp, Idfm, and Idfa). For the prupose of this study, the caudal end of Idfp was grouped with Igd and Igv into a posterior Ig (IgP) and the rest of Idfp and Idfm were grouped into a dorsal Ig (IgD) (**Fig. 4A**, slice 2 showing the boundary of IgP and IgD). Idfa was grouped with Ia (see below). Id contains four distinct dorsoventrally stacked areas: dorsal (Idd), mound (Idm), ventral (Idv) and ventral fundal posterior (Ivfp). We simplified this parcellation, with Idd being here our dorsal Id (IdD) and the other three areas forming our ventral Id (IdV) (**Fig. 4A**, red line showing the boundary of IdD and IdV).

Cytoarchitectonically, Ia is located ventral to the fundus. However, functionally, the anterior section of the dorsal fundus is often associated with a greater dorsal anterior insula and was thus included in a general Ia for the purpose of this study. All these procedures are further evaluated by our co-author Henry Evrard (insula expert^48,71^). The cingulate and insula segmentation results were also transformed from volume space to surface space.

Since the amygdala is located within deep brain nuclei and is relatively small in size, automated segmentation using the D99 subcortical atlas alone is less reliable. Therefore, we performed manual-guided segmentation based on the D99 brain atlas (Saleem and Logothetis, 2012).

Specifically, we selected the high-SNR structural image from our dataset that most closely matched the structural image in the atlas, and used the boundaries of the amygdala subregions defined in the D99 atlas to manually align and map them onto our high-resolution structural image (bottom of **Fig. 1D**). This process provided accurate segmentation of the cingulate insula and amygdala, and also further evaluated by our collaborator Katalin Gothard (amygdala expert^57,72^).

### Functional MRI data preprocessing

The EPI functional images were preprocessed with distortion correction, slice timing correction and motion correction. Images across time points were averaged after motion correction. Then this averaged functional image was registered to the high SNR anatomical image using Boundary-Based Registration (BBR)^106^ to estimate spatial transformation from functional image to anatomical image. The baseline drift and high-frequency components were further removed from motion corrected data with a third-order polynomial function and a bandpass filter of 0.01-0.08 Hz^107^. The cerebrospinal fluid (CSF) signal and global signal were extracted from the CSF mask and whole-brain mask, both derived from the surface-reconstruction segmentation, for further regression analysis.

### fMRI statistical analysis

For each stimulation site, timecourses from all experimental runs (15 trials per run) at the intensity of 0.3 J/cm^2^ were included in the statistical analysis (Monkey P: 45, 30, and 45 trials for sites P1-P3; Monkey Y: 30, 30, and 45 trials for sites Y1-Y3, respectively). A General Linear Model (GLM) analysis was then performed using AFNI to evaluate the significant level of activation induced by INS. The timecourse for each voxel was modeled as a linear combination of predictor variables, including INS stimulus, cerebrospinal fluid (CSF) signal, the global signal, and a residual. The INS stimulus predictor was the convolution of the laser onsets and a canonical hemodynamic response function (HRF). For each voxel, the beta weights for the INS stimulus predictor were subjected to student’s T-test to determine if the beta weights were significantly different from zero. Significance values were FDR-corrected (Benjamini-Hochberg) for multiple comparison correction.

### Evaluation of reliability

To demonstrate the reliability of these connections: (1) We qualitatively compared the INS-evoked activation patterns with known anatomical connections^21^ (**Fig. 2**) and found that they occurred in similar locations. (2) The reproducibility of INS-evoked connections was assessed by dividing all experimental trials into independent odd and even subsets (**Fig. S1**, 22 even and 23 odd trials, for a total of 45 trials). Statistical analyses performed on each subset yielded consistent activation patterns in the cingulate, insula, and amygdala (white arrowheads in **Fig. S1**). (3) Maps with different statistically significant level (different p-values, FDR-corrected p = 0.005, 0.01, 0.05, and unthresholded) were also demonstrated to show the stability of the spatial trend of activations at different thresholds (**Fig. S2**). (4) The intensity dependence test was also performed to show the relationship between the signal amplitudes and INS intensities for the time series extracted around the fiber tip (**Fig. 1H**).

### Evaluation of patch size and overlay index

To quantify the spatial characterization of the connected patches, we calculated the area of each patch in cingulate and insular cortex, using *mris_anatomical_stats* in Freesurfer. Assuming the activation patches were circular, we estimated their approximate diameters using the formula (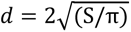) for the area of a circle. To examine the similarity of spatial location between these patches, the Dice coefficient was used to quantify the overlap ratio. Specifically, the number of intersected vertices was divided by the total number of the activated vertices. For amygdala activations, the size of an activation cluster was calculated by multiplying the number of voxels by the voxel size (each cluster was defined by face-connected voxels).

### Evaluation of connectivity topography

For each of the three PM stimulation sites in each monkey, activation patches (activated voxels) were color-coded in red (site P1 and Y1), green (site P2 and Y2) and blue (site P3 and Y3).

These patches (voxels) were then outlined to examine their spatial distribution within each target brain region (cingulate, insula and amygdala) and each subregion (four anterior-to-posterior groups in area 24^67^; three dorsal-to-ventral insular subdivisions in IgD and Id^48^; and each amygdala subdivision BA, LA, AB, CE/AAA). To further compare connectivity topography across all six PM stimulation sites from the two monkeys, we conducted additional analysis, we aligned the stimulation sites and their corresponding activation patterns, to assess the potential shifts or clustering of the activation (see **Figs. S4-S6**).

### Histological procedures

After finishing all MRI sessions, Monkey P was given an overdose of sodium pentobarbital (> 50 mg/kg body weight, IV) and was perfused through the heart with sucrose solution. Weil’s Iron–Hematein method was used for nerve fiber staining in the pulvinar^108^. The detailed procedures, as well as the identification of borders across PM, PL, and PI were described in our previous study^56^. The fiber tract from the last experiment (a PL stimulation experiment) was used as a landmark to align Nissl-stained section and MRI anatomical image that acquired for fiber track determination. This anatomical image was further aligned to the high SNR anatomical image. As a consequence, the staining results were able to overlayed on the high SNR anatomical images for visualization purpose (**Fig. 1C**).

---

**Fig. S1.**
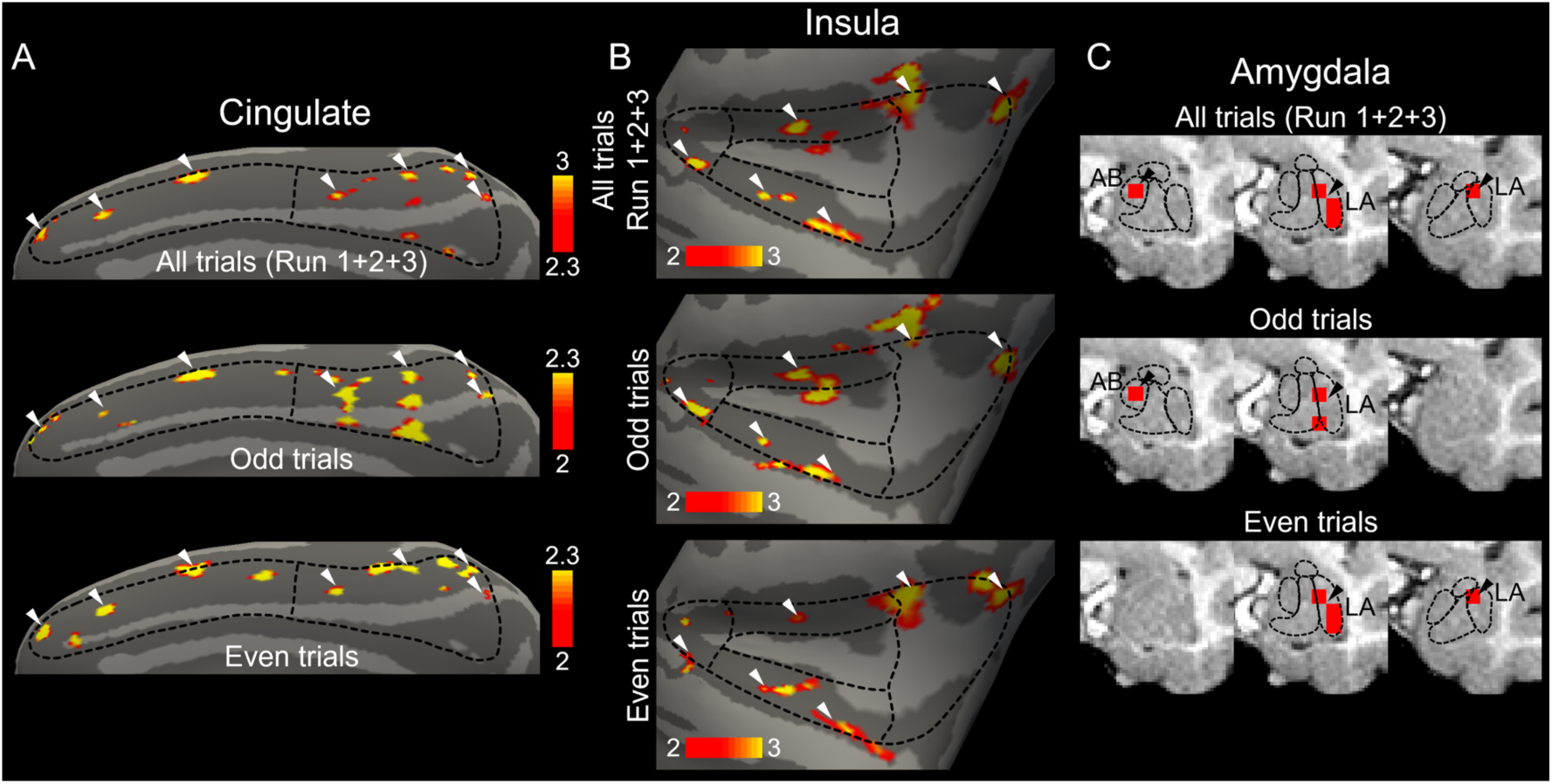
Reproducibility across trials. White arrowheads highlight consistent activation patches across different trials (half-half analysis). (**A-C**) Sums of different trials (all trials; odd trials; even trials) reveal similar cingulate (**A**, FDR-corrected P<0.005), insula (**B**, FDR-corrected P<0.01) and amygdala (**C**, FDR-corrected P<0.005) activation maps. All panels are from monkey P stimulation P1, color bar threshold: -log(p).

**Fig. S2.**
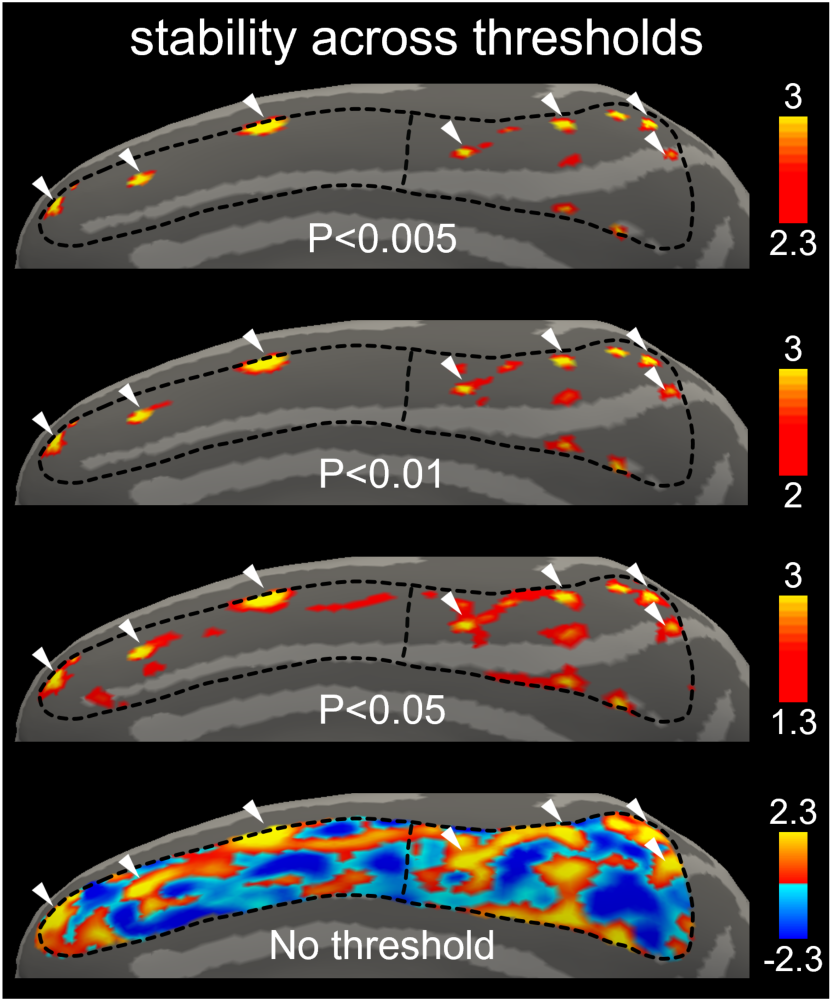
Stability across thresholds. White arrowheads highlight consistent activation patches across different thresholds. Comparison of cingulate activations with thresholds at p<0.005, P<0.01, P<0.05 (all FDR-corrected) and no-threshold show that, while the sizes of patches increase with lower threshold, the activation patterns remain largely stable. All panels are from monkey P stimulation P1, color bar threshold: -log(p).

**Fig. S3.**
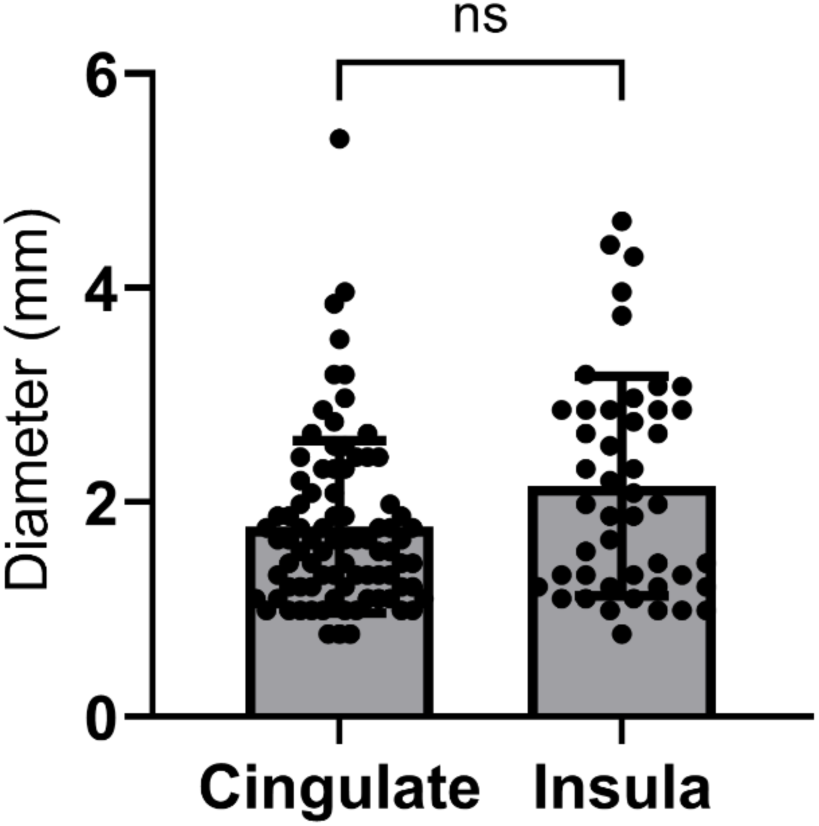
Statistical comparison of patch diameters in cingulate and insula. A nonparametric Mann–Whitney U test revealed no significant difference in patch diameter between the cingulate cortex (2 cases, n = 85) and the insula (2 cases, n = 46).

**Fig. S4.**
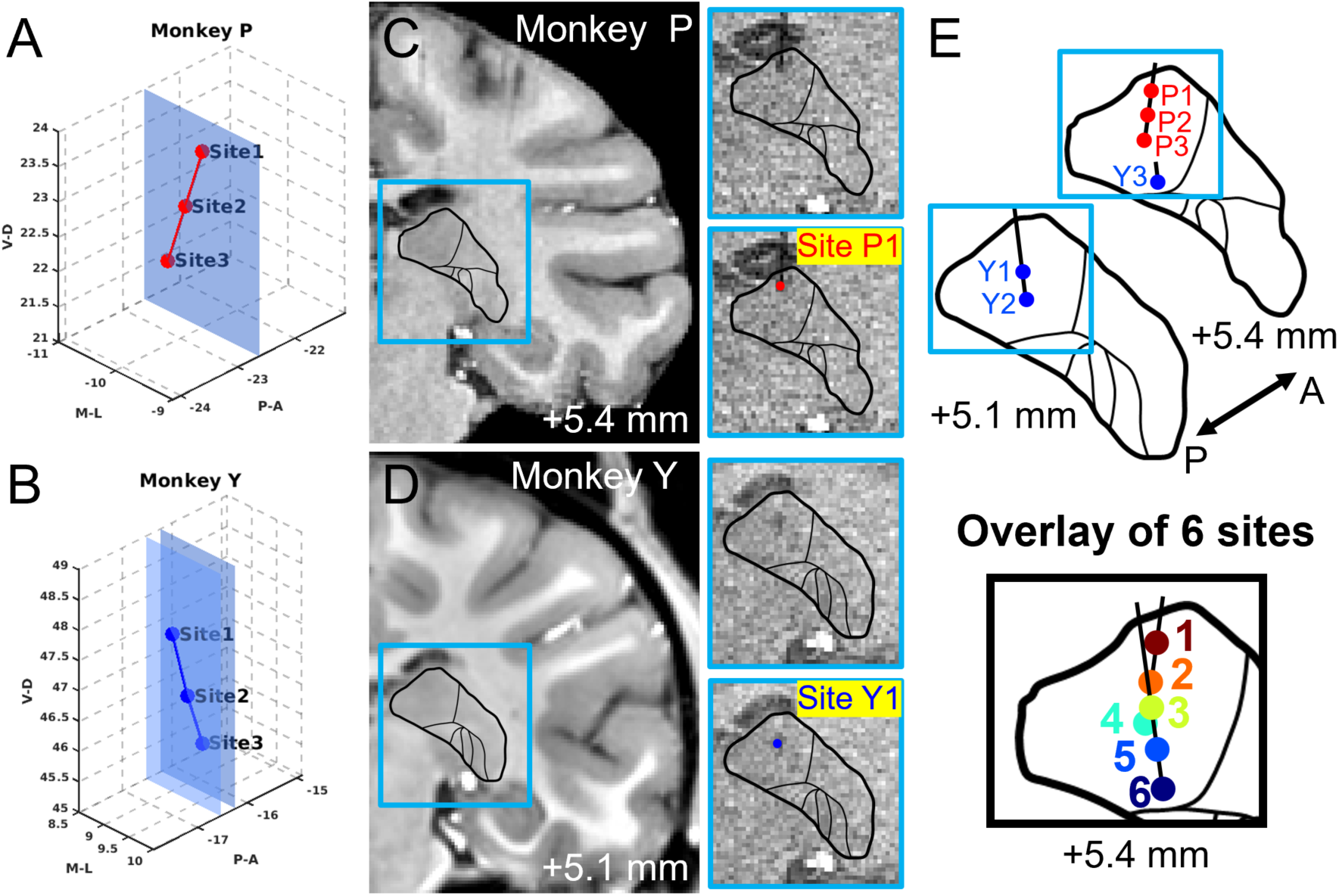
Identification and combination of stimulation sites in 2 monkeys. The combined positions of different PM stimulation sites in two monkeys based on stereotaxic coordinates and anatomical staining. (**A-B**) The three stimulation sites in monkey P (**A**) are located within the same coronal slice (+5.4 mm from the ear bar), while in monkey Y (**B**), stimulation sites 1 and 2 lie within one coronal slice (+5.1 mm from the ear bar), and site 3 lies in a different coronal slice (+5.4 mm from the ear bar). (**C-D**) By aligning the boundary of the pulvinar (in the blue box) obtained from anatomical staining, spatial alignment of the pulvinar across the two animals were achieved. The left panels display the alignment with the high-SNR structural image in monkey P (**C**) and Monkey Y (**D**), and the right panels show the raw MPRAGE structural image at the same location, along with the stimulation sites identified (Monkey P: red dot in **C**, Monkey Y: blue dot in **D**). (**E**) Given the close proximity of the slices across the two monkeys (top panel, red dots: sites P1, P2 and P3 in Monkey P, blue dots: sites Y1, Y2 and Y3 in Monkey P, two adjacent slices), these slices were collapsed onto a single representative slice (bottom panel, +5.4 mm from the ear bar), the combined sites 1-6 correspond to P1, P2, Y1, P3, Y2, and Y3, respectively.

**Fig. S5.**
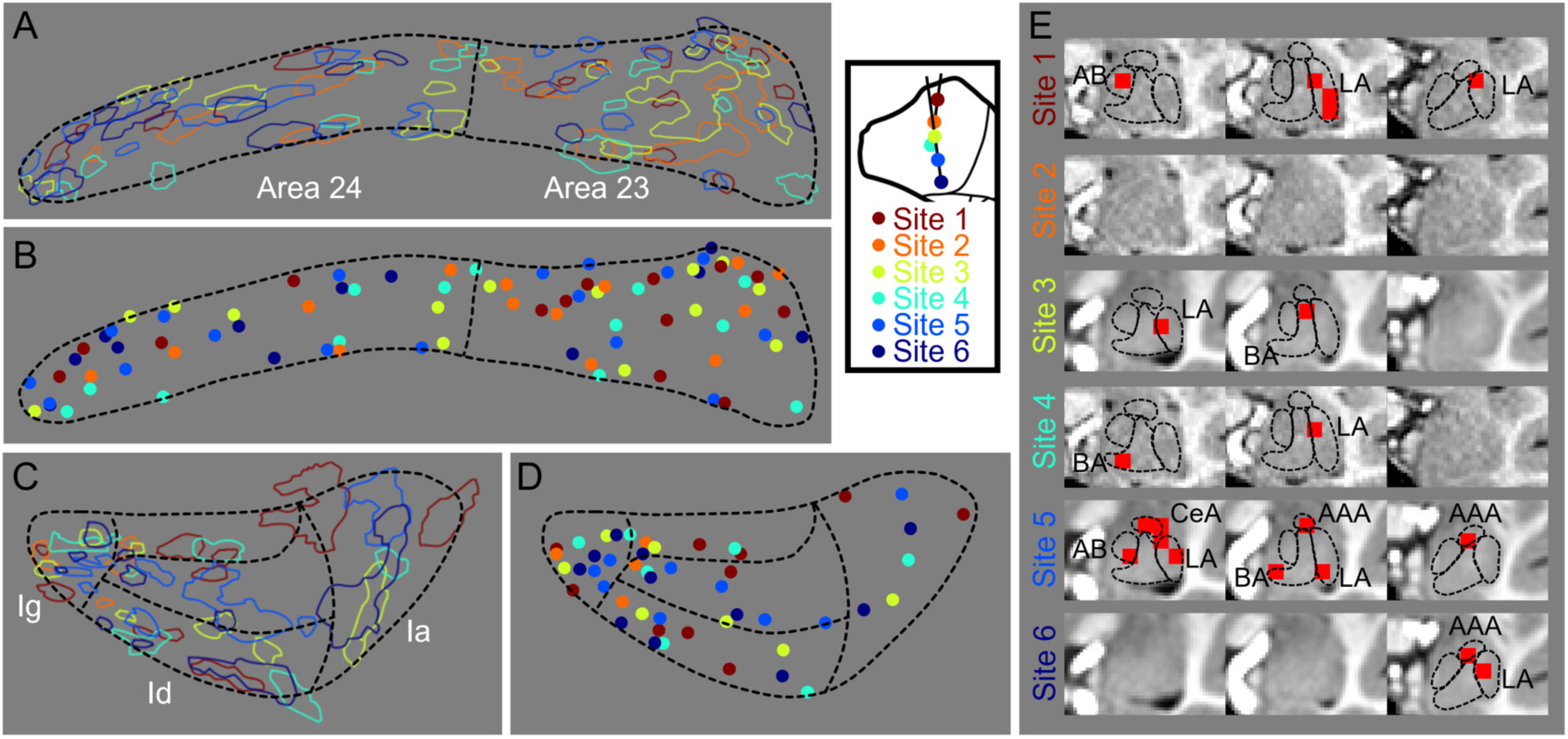
Activation maps of six sequential sites after alignment. No systematic order was found. (**A-D**) To account for minor size differences between the two monkeys, the brain surface maps were slightly rescaled based on the boundaries of the cingulate and insula. Activated patches were then overlaid and color-coded according to the sequence shown above. Combined activation maps are shown for the cingulate (**A**) and insula (**C**). For visualization of patch locations, each activated patch is additionally represented by a single dot, defined by the surface vertex with the highest t-value within that patch, in the cingulate (**B**) and insula (**D**). (**E**) The sequential activation maps of the amygdala (sites 1-6) are also shown.

**Fig. S6.**
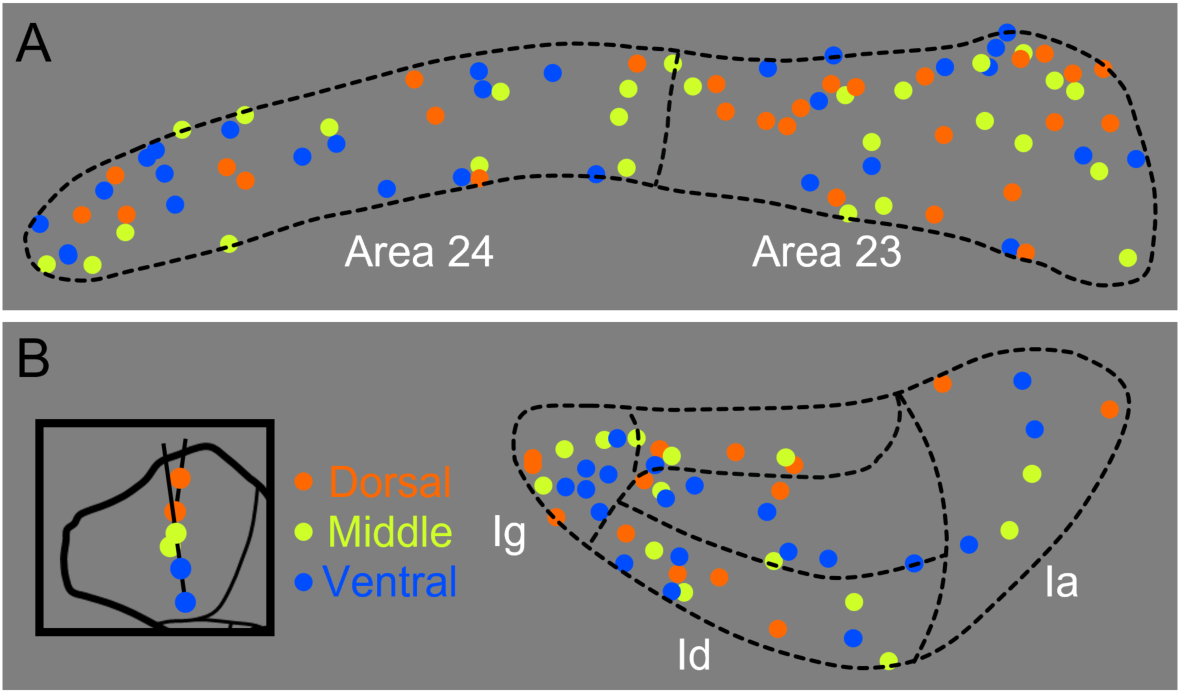
Activation maps of dorsal/middle/ventral PM stimulation sites. No systematic order was found. (**A-B**) The six PM stimulation sites were also classified into dorsal (orange), middle (yellow), and ventral (blue) categories based on their relative positions along the dorsoventral axis. Patch centers were color-coded accordingly in the cingulate (**A**) and insula (**B**).

